# Antibodies to influenza A virus hemagglutinin and neuraminidase limit egress and alter the physical properties of released virus particles

**DOI:** 10.64898/2026.06.15.729605

**Authors:** Anna Jaeggi-Wong, Antonio Santos-Peral, Edward A. Partlow, Tongyu Liu, Tijana Ivanovic

## Abstract

Influenza A virus (IAV)-specific antibodies neutralize mature virions by inhibiting functional sites or, in some cases, by promoting virion aggregation. Many antibodies also act on infected cells to reduce virion yields, but the underlying mechanisms and effects on the physical properties and function of released virus particles remain incompletely characterized. Here we use flow virometry to acquire high-sensitivity yield and size measurements of virus particles released in the presence of antibodies. Combined with digital-droplet PCR and electron microscopy, this approach enables comprehensive characterization of antibody-induced changes in particle genome-content and morphology. We show that antibodies rapidly and dynamically alter released particle distributions, reducing yields and inducing the production of larger particles. Both effects result in part from aggregation induced by crosslinking of viral antigen on the infected-cell surface and inhibition of viral NA. However, yield reduction is not fully explained by aggregation, and a subset of induced larger particles are elongated virions. Finally, particles formed in the presence of HA stem-binding antibody, which does not inhibit attachment of mature virions, show reduced attachment in subsequent rounds of infection. Altogether, we uncover an unappreciated mechanism by which antibodies interfere with viral infection that occurs only during budding and release. Our work highlights the necessity of studying how the immune response shapes virus populations in the context of active infection processes.

## Introduction

Viruses and their human hosts have coevolved throughout evolutionary history. Humans have developed immune responses to control virus infection, which in turn impose evolutionary pressures that shape virus populations through direct effects on free virions or infected cells or indirect effects on bystander cells. Understanding these virus-host interactions underlies the development of more effective vaccines and therapeutics. Despite this, there is limited understanding of how immune pressures, such as antiviral antibodies, affect the physical properties of virus particles, and whether these changes inhibit or facilitate downstream infection. Influenza A virus (IAV) is an enveloped virus that assembles at the host plasma membrane, producing particles with a broad size distribution, ranging from about 100 nm spherical virions to filamentous forms tens of microns in length (1, 2). During infection, the newly produced transmembrane viral surface glycoproteins hemagglutinin (HA) and neuraminidase (NA) localize to lipid rafts on the cell surface (3). The matrix protein 1 oligomerizes beneath the membrane, inducing budding and recruiting the viral genome (4, 5). The matrix protein 2 (M2) is implicated in membrane scission (6), and NA facilitates virion release from the infected-cell surface by cleaving interactions between HA and sialylated cell-surface receptors (7). Although HA, NA, and M2 are all abundant on the infected-cell surface (8, 9), their incorporation into virions is markedly asymmetric. HA covers about 80% and NA about 20% of the virion surface, and M2 is present in relatively few copies (10, 11). As a result, these proteins represent key, but variably displayed, targets for antibody responses, with HA dominating natural responses and NA and M2 being less immunogenic but more conserved.

Antibodies to HA typically neutralize virions, inhibiting cell entry. Antibodies targeting the HA head domain block virion attachment to host cell receptors (12–14), while antibodies to the HA stem inhibit fusion of the viral and host membranes after endocytosis (15–20). In addition to these canonical antibody functions, some HA head-binding bivalent IgGs (21) and dimeric IgAs (22) were shown to inhibit infection by promoting virion aggregation, effectively reducing the number of independent infectious units. While HA stem-binding IgGs alone do not block attachment or induce virion aggregation (21), they were shown to inhibit attachment in the presence of the multivalent complement component C1q (23), revealing that antibody effects can vary depending on the environment in which they act.

In addition to neutralizing free virions, antibodies can act on infected cells through Fc-mediated effector functions. Binding of HA stem, NA, and M2 antibodies to antigens on the surface of infected cells can trigger their removal or killing by antibody-dependent cellular phagocytosis, antibody-dependent cellular cytotoxicity, and complement-dependent cytotoxicity (24–31). In the absence of cell killing, antibodies against all three viral surface glycoproteins were shown to reduce infectious virion yields (11, 12, 22, 32–37), and some mechanisms have been identified. For example, some HA and NA antibodies inhibit virion release by inhibiting NA activity, either through direct binding of NA antibodies to the active site (38) or through steric hindrance by the Fc region of HA-bound antibodies (35, 39, 40). Other HA antibodies can promote particle retention by crosslinking HA on the infected cell surface, either within the membrane or between the cell and virions (12, 22, 32). Finally, M2-targeting antibodies have been associated with reduced surface M2 expression (33).

Emerging evidence suggests that perturbations of budding and release can affect not only yield but also the properties of released particles, including composition, size, and shape. Virions produced in the presence of the small molecule NA inhibitor oseltamivir had a higher NA to HA ratio and were shorter in length than those produced in the absence of treatment (41). Furthermore, treatment of infected cells with a large panel of antibodies targeting all three IAV surface proteins across multiple epitopes and strains shifted the size distributions of released particles toward larger sizes, indicating that changes in released particle properties are not limited to small-molecule perturbations but extend broadly across antibodies (36). Finally, infections by a filamentous IAV strain in the presence of M2 antibodies produced a greater proportion of spherical particles, interpreted to result from virion fragmentation (34). Some of these changes have been proposed to be proviral, others antiviral. For example, under NA inhibition (NAI), the release of virions with higher NA-content might provide replicative advantage (41). Similarly, filamentous virion production in the presence of antibody might be beneficial, as larger particles can resist antibody-mediated inhibition of cell entry (42, 43). Conversely, as IAV filaments package a single genome like spherical particles (44), their fragmentation might be antiviral because it would generate smaller, mostly genome-less particles (45). While antibodies can reduce virion production and alter released virion properties, the mechanisms and resulting changes in virion populations are incompletely characterized, and their links to downstream infection remain limited.

In this study, we identify and characterize novel antiviral effects of antibodies that occur in association with active virus budding and release. We reveal that antibody binding to viral surface antigens on infected cells rapidly and dynamically reduces the yield and alters the physical properties of released virus particles, leading mostly to aggregation, but also to filament formation. We also link these outcomes to antibody-mediated molecular events on the surface of infected cells. Notably, aggregation has both NAI-dependent and independent causes, and some yield reduction occurs independently of aggregation. Finally, although HA stem antibody binding to free mature virions does not interfere with particle attachment in subsequent infections (23), we find its binding during particle production does, propagating antibody effects beyond the initial infection. Altogether, this work highlights the necessity of studying how virus-host interactions shape not only cellular infection outcomes but also the properties of virus particles, and the importance of doing so in the context of active infection processes.

## Results

### Antibodies to major viral surface glycoproteins rapidly alter virus particle production

To characterize the short-term effects of antibodies on virus particle production, we infected cells with Puerto Rico/8/1934 IAV (PR8) at multiplicity of infection (MOI) 0.6 and treated them with antibodies targeting the HA stem (MEDI8852 IgG) or NA (NA2-1C1 IgG and NA2-10E10 IgG) from 21 to 23 hours post infection (h.p.i.). We sampled the cell supernatants at 15- or 30-minute intervals during antibody treatment and performed flow virometry (36) to count virus particles and measure their size distributions (Fig. 1A). Virion counts deriving from mock-treated cells steadily increased, and particle size distributions remained stable (Fig. 1B-C), as expected at this time post-infection (36). In contrast, HA stem antibody reduced released particle yield and increased the fraction of large particles within 15 minutes of treatment (Fig. 1B). Notably, the extent of large particle induction by HA stem antibody was not monotonic. The initial rise waned and was followed by a second size shift that emerged by 120 minutes post-treatment, revealing an active dynamic between antibodies and infected cells. NA antibodies resulted in similar effects on yields, greater magnitude effects on size, and less pronounced size dynamics (Fig. 1C).

**Figure 1.**
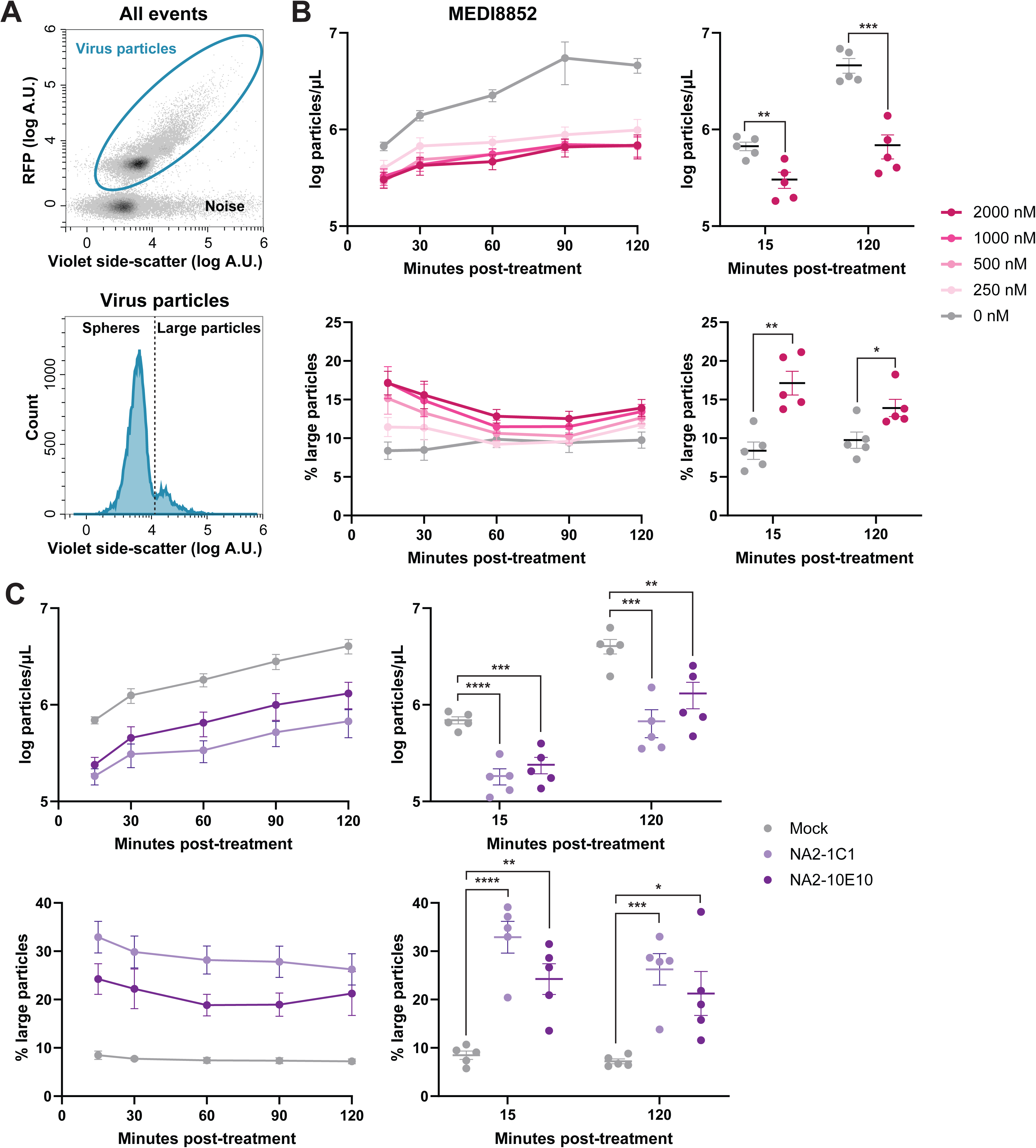
Antibodies to major viral surface glycoproteins rapidly reduce yield and alter virus particle size distributions. **A)** Flow virometry for quantification of virus particle yield and size. Virus particles are separated from noise by labeling with a fluorescent antibody targeting HA (top). Spherical and large particle fractions are distinguished by violet side-scatter (bottom). **B-C)** MDCK cells infected with PR8 at MOI 0.6 for 21 hours were washed into media containing **B)** the indicated concentrations of MEDI8852 IgG or **C)** 500 nM of the indicated NA antibody. A no-antibody, mock-treated sample was included in each experiment. Virus particle yield (top) and size (bottom) from the infected-cell supernatant at the indicated times (left). Yield and size for the highest antibody concentration (right) at 15- and 120-minutes post-treatment. Supernatants were analyzed by flow virometry as in A. Mean and S.E.M. are plotted for *N*=5 biological replicates. *P* values shown are from two-sided unpaired *t*-tests. \**P*<0.05; \*\**P*<0.01; \*\*\**P*<0.001; \*\*\*\**P*<0.0001.

While flow virometry enables fast and sensitive measurements, it does not readily distinguish single larger virions from aggregates of multiple smaller virions. We thus hypothesized that antibodies might be inducing virion filamentation, causing their aggregation, or exerting a combination of these effects. To test whether the treatment antibodies can crosslink virus particles post-release, we incubated the mock-treated supernatants with the HA stem or NA antibody at the highest concentration and under the same conditions used in the budding experiments. Neither antibody reduced particle counts or increased the proportion of large particles, indicating that neither antibody aggregates mature virions under our assay conditions (Fig. S1).

We also tested M2-targeting antibodies 14C2 and monoclonal antibody 65 (mAb65). Neither antibody affected particle yields in the budding experiment, and although they caused a slight increase in the fraction of large particles (Fig. S2A), they also aggregated free virions to equivalent extents (Fig. S2B). Together, these results show that HA and NA antibodies, but not M2 antibodies, rapidly reduce particle yields. They all dynamically alter particle size distributions to various extents during budding, but only M2-antibody effects can be explained by post-release crosslinking.

### Antigen crosslinking contributes to the initial changes in IAV distributions

Since crosslinking by bivalent IgGs can contribute to their activity, we probed the contribution of antibody valency and size to changes in IAV yield and particle-size distributions. We performed the short-term treatments of PR8-infected cells with a panel of HA stem antibody fragments. F(ab’)_2_ fragments, which lack the Fc domain but retain both antigen-binding arms, recapitulated the particle counts and size distributions induced by the full-length IgG with the same dynamics (Fig. 2A). This revealed that the antigen binding arms, without steric occlusion of NA by the Fc region, were sufficient to induce the observed HA stem-antibody effects. We next tested smaller, monovalent antibody fragments. Neither Fab nor single chain (scFv) fragments reduced particle count (Fig. 2A) despite binding to HA (Fig. S3). In fact, the Fab fragments increased yield relative to the mock-treated condition (Fig. 2A). Additionally, neither fragment increased larger-size particle fraction at the earliest timepoints (Fig. 2A). These results suggest that crosslinking of antigen by bivalent antibodies is necessary to reduce yield and induce the initial size shift. Notably, the Fab fragment produced the second size shift with similar magnitude and dynamics as IgG and F(ab’)_2_ (Fig. 2A), showing that the delayed size-shift is crosslinking-independent. Different dependencies of early and late size shifts suggest they are governed by different underlying mechanisms, while yield reduction and the initial rise or decay of the large-particle fraction might be functionally linked.

**Figure 2.**
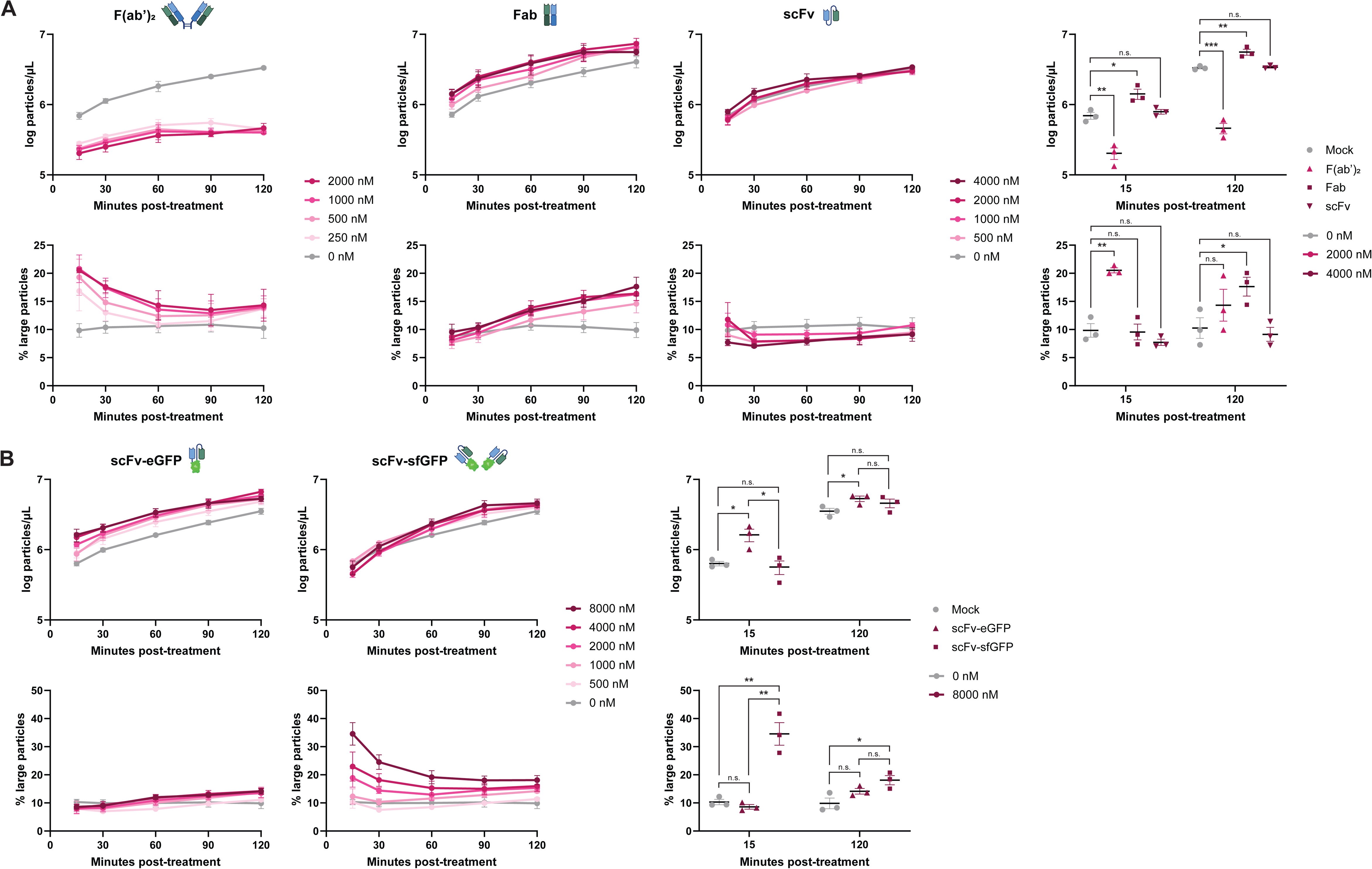
Antigen crosslinking is required for the initial changes in IAV distributions. MDCK cells infected with PR8 at MOI 0.6 for 21 hours were washed into media containing the indicated concentrations of **A)** MEDI8852 F(ab’)_2_, Fab, or single-chain (scFv) fragments or **B)** MEDI8852 single-chain (scFv) fragment tagged with monomeric enhanced green fluorescent protein (eGFP) or weak dimer superfolder GFP (sfGFP). A no-antibody, mock-treated sample was included in each experiment. Virus particle yield (top) and size (bottom) from the infected-cell supernatant at indicated times (left). Supernatants were analyzed by flow virometry. Yield and size for 2000 nM F(ab’)_2_ and 4000 nM Fab and scFv (in panel A) and 8000 nM scFv-eGFP and scFv-sfGFP (in panel B) at 15- and 120-minutes post-treatment (right). Mean and S.E.M. are plotted for *N*=3 (all except Fab<4000 nM) or *N*=4 (Fab<4000 nM) biological replicates. *P* values shown are from two-sided unpaired *t*-tests. n.s., not significant, *P*>0.05; \**P*<0.05; \*\**P*<0.01; \*\*\**P*<0.001. Icons created in BioRender. Ivanovic, T. (2026) https://BioRender.com/u7cy3sg

To directly test the contribution of antibody valency to the yield or early time-point size shifts without confounding differences in construct size, we fused the HA stem single-chain fragment to either superfolder green fluorescent protein (sfGFP), which dimerizes at high local concentrations, or enhanced green fluorescent protein (eGFP), which does not dimerize (46). We additionally generated fusion proteins of the same single-chain fragment and tagRFP-T or its non-dimerizing, single-point mutant E205K (47). We then repeated the time-course experiment. Both dimerization-capable constructs and neither monomeric construct triggered the initial rise and decay in large-particle fraction, like IgG and F(ab’)_2_ (Fig. 2B, S4), validating that bivalent binding, rather than authentic antibody structure, is crucial for those effects. Notably, the dimerizing scFv-tagRFP-T construct strongly reduced particle counts, resembling IgG and F(ab’)_2_, but scFv-sfGFP did not (Fig. 2B, S4), indicating that other features of the antibody fragments, such as dimerization affinity or geometry, are important for yield effects. This result further functionally decoupled yield reduction and the initial size shifts. While the monomeric scFv-eGFP construct increased yield like Fab (Fig. 2B), neither monomeric construct altered the virion size distribution after one hour despite having similar size and the same valency as Fab (Fig. 2B), revealing further complexities in antibody-induced molecular events on the infected cell surface. In sum, while both yield reduction and the initial rise and decay in the large-particle fraction depend on antigen crosslinking, yield reduction is more sensitive to dimerization parameters. Among monovalent constructs, only Fab induced the delayed size increase, revealing that these effects are independent of antigen crosslinking but depend on other molecular features not recapitulated in the chimeric constructs. Finally, both Fab and scFv-eGFP increased yield, opposite to the effect of the authentic bivalent IgG. We suggest molecular interpretations to reconcile these observations in the Discussion. These combined experiments reveal complex dynamics, in which yield and size-distribution effects, as well as early and delayed size shifts, can be uncoupled from one another. The complex and rapid antibody-induced changes in particle distributions further underscore the utility of high-sensitivity flow virometry for studying antibody-virus interactions during assembly and release.

### Antibody binding triggers internalization of viral antigen

Many endocytic pathways begin with dimerization of the transmembrane cargo (48, 49). Although viral glycoproteins are not canonical endocytic cargo, we hypothesized that their crosslinking by antibody, which underlies changes in particle distributions, might also drive antigen internalization. We treated PR8-infected cells with HA stem, NA, or M2 antibodies from 21 to 23 h.p.i., then visualized antibody localization by confocal immunofluorescence microscopy. Consistent with prior observations (36), HA stem antibody localized to both the surface and interior of infected cells, forming puncta in the cytoplasm and perinuclear space after treatment (Fig. 3A). Antibody internalization was not limited to HA stem antibody but was also evident for NA (Fig. 3B) and M2 (Fig. 3C) antibodies. Across these antibodies, puncta varied qualitatively and correlated with the magnitude of their effects on virus particle distributions. For example, antibodies targeting the HA stem were more diffuse on the cell surface and formed smaller puncta inside cells, while antibodies targeting NA formed larger puncta both on the surface and in the cytoplasm (Fig. 3A-B). M2 antibodies appeared the most diffuse and formed the smallest puncta (Fig. 3C). This active interplay between antibodies and infected cells might be related to their dynamic effects on particle size distributions. Furthermore, we hypothesized that the antibody-induced internalization might limit the amount of viral antigen on the cell surface available for budding, contributing to the yield reduction and/or altering the structure or organization of released virions.

**Figure 3.**
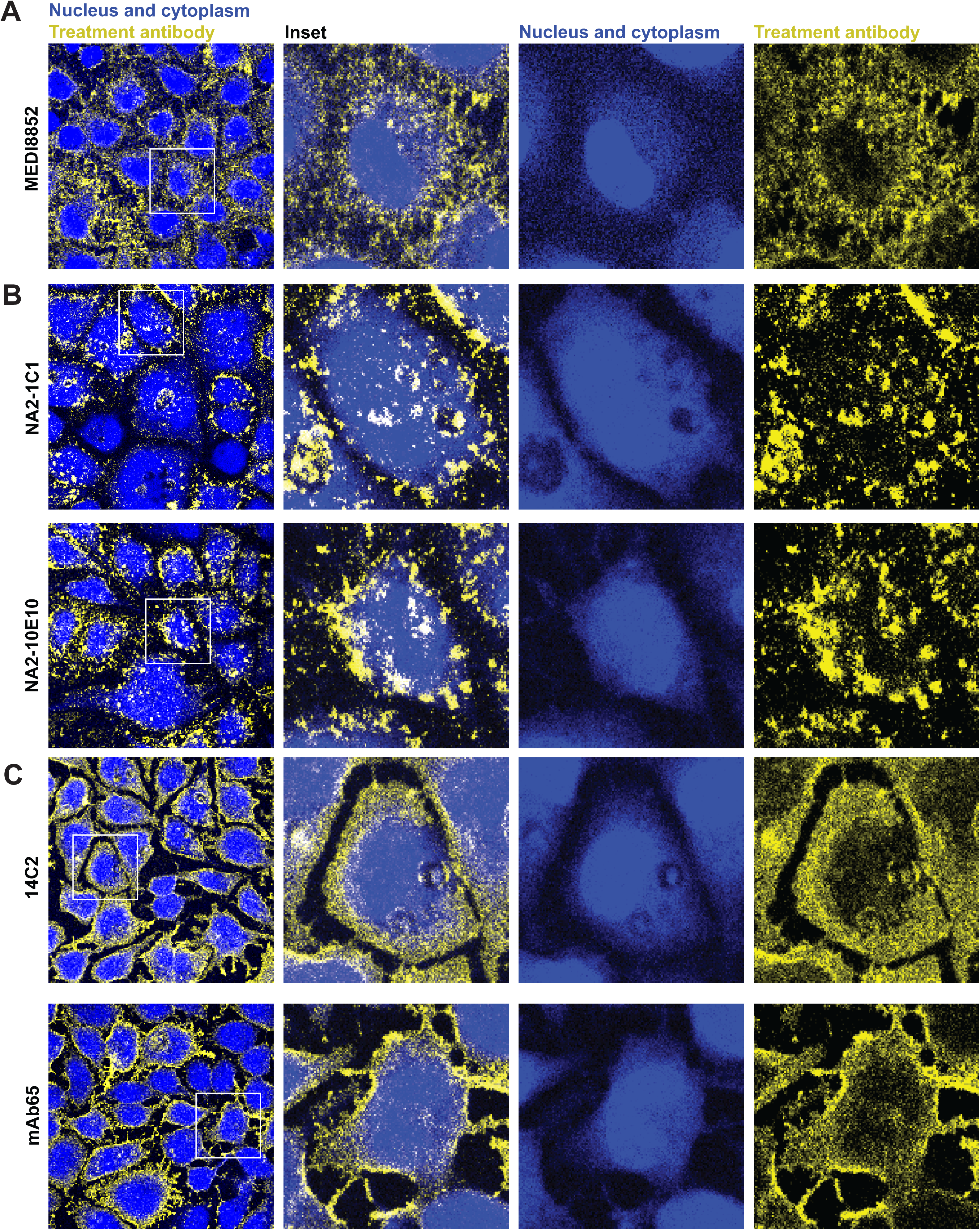
Antibody binding triggers internalization of viral antigen. MDCK cells infected with PR8 and treated at 21 hours post-infection with 500 nM of the indicated **A)** HA stem, **B)** NA, and **C)** M2 antibodies were collected 120 minutes post-treatment and analyzed by immunofluorescence. Representative images showing nuclei and cytoplasm (blue) and total cell-associated antibody treatment (yellow).

### Antibody-induced large particles include both aggregates and filaments

To distinguish whether antibody-induced particle size shifts arose from aggregation or filamentation, we measured average genome content per particle, or ploidy, by combining digital droplet PCR (ddPCR) to count genomes with flow virometry to count virus particles. Because IAV packages at most a single genome-complement per virion (44), increased ploidy would indicate particle aggregation, whereas increased particle size without increased ploidy would support filamentation. We treated PR8-infected cells with HA stem or NA antibodies at 22 h.p.i. and harvested the supernatant at 24 h.p.i for analysis (Fig. 4, *filled circles*). We used oseltamivir, a small-molecule viral NA inhibitor, as a control. It inhibits particle release by promoting sialic acid-mediated retention to cells and association between budding particles (50, 51), and particles released in its presence are thus likely to include aggregates. As expected, oseltamivir reduced particle yield, increased the proportion of large particles, and raised the average ploidy relative to the mock treatment (Fig. 4, S5A, *filled circles*). Both HA stem and NA antibodies also increased ploidy in association with an increase in the large-particle fraction. NA antibody effects were stronger than those of HA stem antibody and comparable to oseltamivir (3 to 8-fold for NA, 2 to 3-fold for HA stem, and 4 to 8-fold for oseltamivir) (Fig. 4, S5A). These results show that NAI by small-molecule drugs and antibodies induce virion aggregation during active budding and release, resulting in fewer, larger particles with increased genome content.

**Figure 4.**
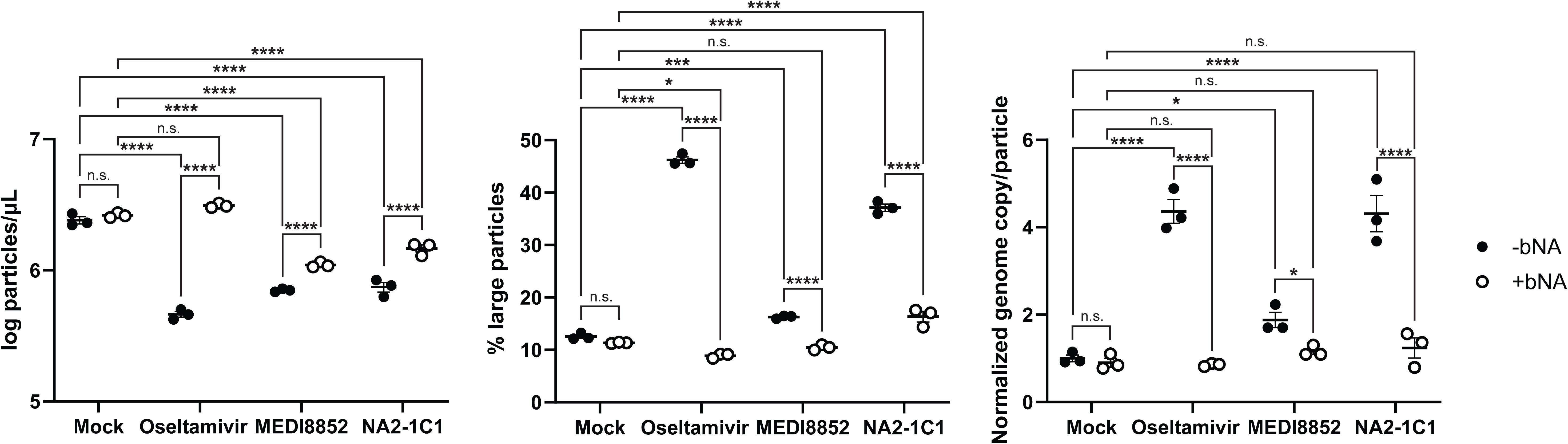
Antibodies increase particle ploidy and produce NAI-mediated aggregates. MDCK cells were infected with PR8 at MOI 0.6 and treated at 22 hours post-infection with 1000 nM oseltamivir, 2000 nM MEDI8852 IgG, or 500 nM NA2-1C1 IgG. Bacterial neuraminidase (bNA) was included in the media for the +bNA condition from 20 to 24 hours post-infection. No-bNA (-bNA) and no-antibody (mock) samples were included. Supernatants collected 2 hours post-treatment were analyzed by flow virometry and digital droplet PCR (ddPCR). Virus particle yield (left), size (center), and genome copy per particle normalized to the no-bNA condition (right) from the infected-cell supernatant. Mean and S.E.M. are plotted for *n*=3 technical replicates. *P* values shown are from ordinary 2-way ANOVA with Tukey’s multiple comparisons test. n.s., not significant, *P*>0.05; \**P*<0.05; \*\*\**P*<0.001; \*\*\*\**P*<0.0001.

To investigate the underlying mechanism and determine if aggregation accounts fully for measured changes in particle size or yields, we explored whether NAI by antibodies contributes to virion aggregation during active budding. We pretreated infected cells with bacterial neuraminidase (bNA) at 20 h.p.i. to deplete sialic acid and abolish the requirement for viral NA activity for virion release. We then treated infections with oseltamivir or antibody in the continued presence of bNA from 22 to 24 h.p.i., as above. bNA pretreatment completely prevented oseltamivir effects, with particle count, large-particle fraction, and ploidy remaining at mock treated levels (Fig. 4, S5A, *open circles*). Similarly, bNA treatment of particles produced in the presence of oseltamivir without bNA pretreatment de-aggregated clumps, completely resolving size changes (Fig. S5B). The combined results with oseltamivir thus validate the use of bNA to negate NAI effects during particle release and to resolve sialic acid-dependent aggregation in released virions. Notably, bNA also almost completely resolved oseltamivir-treatment yields to mock levels (Fig. S5B), indicating that cell surface retention of budding particles is only a minor contributor to yield reduction by oseltamivir. Rather, NAI primarily reduces yields by aggregating released particles.

The consequences of bNA treatments on antibody effects varied and revealed variable contributions of NAI to changes in released particle yield, ploidy, and size (Fig. 4, S5). In experiments with HA stem antibody, bNA pretreatment only slightly recovered particle counts (Fig. 4, S5A), exposing effects on yield beyond NAI. In some experiments, bNA pretreatment completely resolved size shifts and ploidy (Fig. 4), whereas in others, it only slightly decreased large-particle fraction and had no effect on ploidy (Fig. S5A). This demonstrated that NAI contributes to, but is not the only source of, aggregation by HA stem antibodies. In experiments with NA antibody, bNA pretreatment resulted in marked but incomplete recovery of particle counts and larger-particle fraction, along with reversion of ploidy ranging from partial to complete (Fig. 4, S5A). This indicated that aggregation is a major component of NA antibody effects and is in large part NAI-mediated. Since the fraction of large particles remained elevated in NA antibody-treated samples even when ploidy was fully resolved (Fig. 4), some increases in particle size under NA antibody treatment are independent of aggregation and might reflect filamentation. Finally, since particle yields were never fully rescued for either HA stem or NA antibodies (Fig. 4, S5A), both treatments additionally attenuate virion assembly or release by an aggregation-independent mechanism. bNA treatment of released virion supernatants (Fig. S5B) supported the conclusions from bNA pretreatment of cells (Fig. 4, S5A). Together, these results show that HA stem and NA antibodies cause aggregation of released virus particles in part by NAI. This aggregation contributes to but does not fully account for reduced virion yields. Finally, some size shifts arise from NAI-independent aggregation or an increase in size of individual virions.

To directly visualize the morphology of released particles, we performed negative-stain electron microscopy (EM) on purified, concentrated samples from large-scale PR8 infections treated with HA stem or NA antibodies from 4 to 24 h.p.i.. These long-term treatments produced high enough yields to perform EM, while maintaining similar yield and size trends as short-term (120 minute) treatment (Fig. S6). Both mock and HA stem antibody samples contained a high proportion of small particles, 110-150 nm in length, but mock had more (71% and 53% of particles, respectively) (Fig. 5A, *image a and b*, 5B). The HA stem antibody samples contained a greater proportion of filamentous particles over 150 nm (34%) compared to the mock condition (24%) and were enriched for filaments around 300 nm in length (Fig. 5A, *image c*, 5B). In addition, HA stem samples uniquely contained larger, amorphous particles that were not a distinct spherical or filamentous morphology (Fig. 5A, *image d*). There also appeared to be more aggregates containing larger, aberrant particles in the HA stem samples (Fig. 5A, *image e*). Clumped amorphous or aberrant particles were not included in quantification, as we could not accurately measure the length of overlapping particles. Regardless, the genuine filaments and larger amorphous particles revealed by EM were only a minor population. In contrast to the extended filaments induced by HA stem antibody, the predominant population in NA antibody samples was shifted towards slightly elongated virions 150-170 nm in length (Fig. 5A, *image f*), accounting for 24% of virions in the NA antibody condition, compared to 9.2% in mock treated samples (Fig. 6B). These combined EM results reveal antibody-induced enlargement of virions, including filament formation. Since actual virion size shifts are relatively infrequent, EM supports conclusions from virometry and ddPCR that a large fraction of the observed size-distribution shifts derives from aggregates. Altogether, these complementary methods establish that antibodies induce aggregation of released virus particles, which is partially caused by NAI, while also increasing virion size and reducing virion production by aggregation-independent mechanisms.

**Figure 5.**
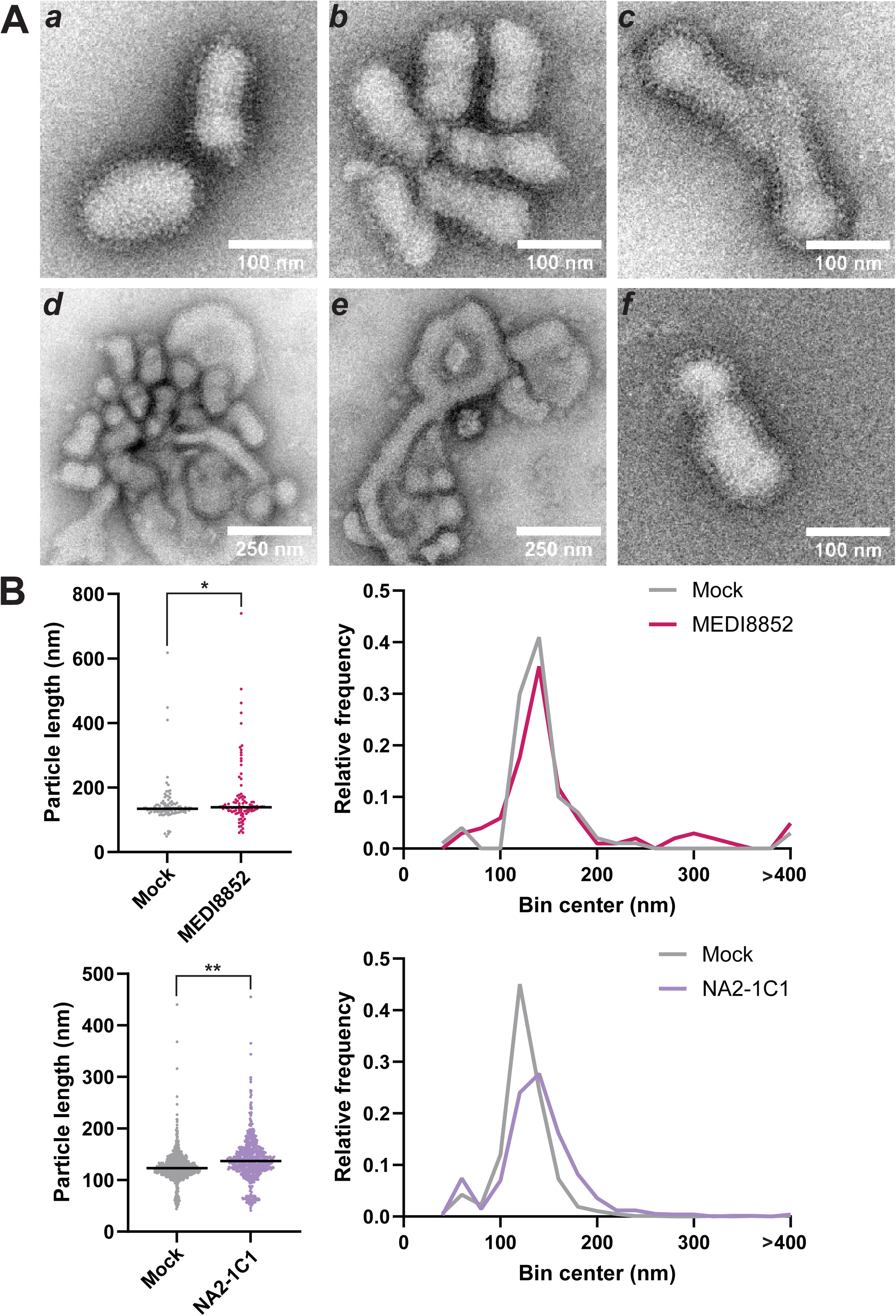
Antibodies induce filamentation. MDCK cells were infected with PR8 at MOI 0.6, treated with 500 nM MEDI8852 IgG or NA2-1C1 IgG or mock treated at 4 hours post-infection, and harvested at 24 hours post-infection. Supernatants were treated with bacterial neuraminidase prior to grid preparation. **A)** Representative EM images for mock (*a*), MEDI8852 IgG (*b-e*), and NA2-1C1 IgG (*f*) treated conditions. **B)** Quantification of particle length from EM and histograms of particle lengths. Bin width=20 nm. All particles greater than 400 nm included in final bin. Median for *n*=100 (mock, left), 116 (MEDI8852 IgG), 852 (mock, right), and 758 (NA2-1C1 IgG) virus particles per condition is plotted. *P* values shown are from one-way ANOVA testing. \**P*<0.05; \*\**P*<0.01.

**Figure 6.**
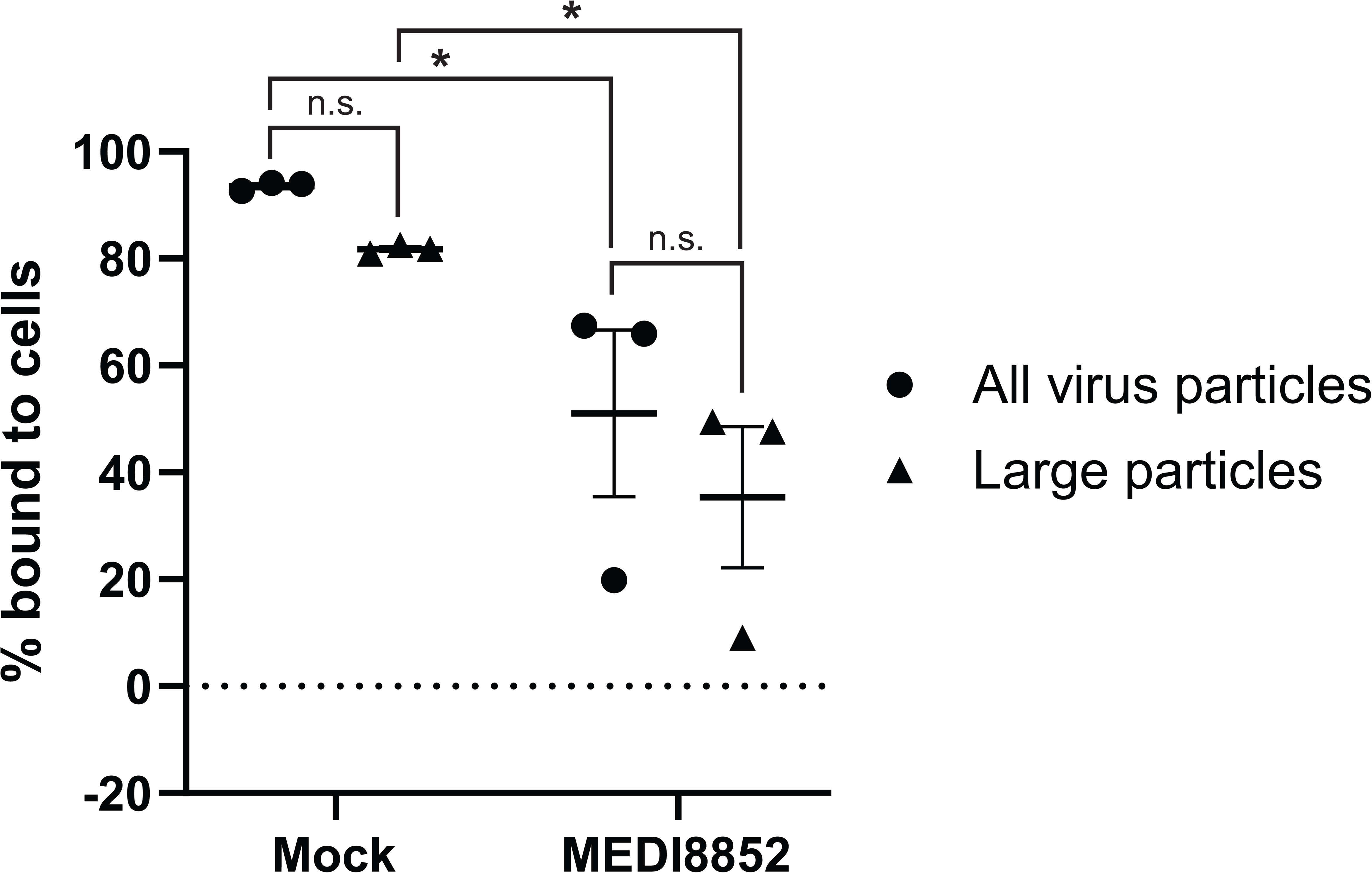
Antibodies inhibit attachment in subsequent rounds of infection. Supernatants harvested from MOI 0.6 PR8 infections after 4 hours of 2000 nM MEDI8852 IgG or mock treatment were incubated for 1 hour at 34°C on MDCK cells. Particles remaining unbound after the attachment period were collected. Inputs and unbound particles were analyzed by flow virometry. Percentage of total input virus or large particles only bound to cells after attachment period. Mean and S.E.M. are plotted for *n*=3 technical replicates. *P* values shown are from 2-way ANOVA Fisher’s LSD test. n.s., not significant, *P*>0.05; \**P*<0.05.

### Antibodies inhibit attachment in subsequent rounds of infection

While the reduction in particle counts from aggregation is antiviral, particle-size increase due to aggregation might be proviral, analogous to how larger, filamentous virions can resist attachment and entry inhibition by antibodies (42). To test whether aggregation promotes or inhibits attachment in new rounds of infection, we harvested virus released from PR8-infected cells after 4 hours of HA stem antibody or mock treatment and incubated it with fresh cells for 1 hour. To quantify binding, we quantified the input virus particles and those remaining unbound after the 1-hour attachment by virometry. We found that virus particles produced in the absence of antibody bound almost completely to fresh cells (93.6 +/- 0.5%), while those produced under antibody pressure had significantly reduced binding (51.1 +/- 15.6%) (Fig. 6). Inhibition of binding in the antibody-treated condition persisted even when quantifying the extent of large-particle binding (81.8 +/- 0.5% for mock versus 35.4 +/- 13.2% for antibody-treated) (Fig. 6, *triangles*), demonstrating that aggregates do not offer a benefit in the attachment step despite their larger size. Virus particles produced in the presence of HA stem antibody are thus neutralized in subsequent rounds of infection despite the same antibody not blocking attachment of free virions.

## Discussion

In this study, we reveal that antibodies present during IAV budding and release dynamically alter released virus particle populations. Antibodies targeting the viral surface antigens HA and NA rapidly reduce particle yields and induce larger-size particles, including NAI-induced aggregates in varying proportions. Antibody-induced effects are complex and multifaceted: the extent of yield reduction or particle size increase and the nature of the large particles vary with the duration of antibody treatment, the target antigen, and antibody valency. We further demonstrate novel, context-dependent antiviral functions of antibodies, including the internalization of antibody-bound viral antigen and attachment inhibition in subsequent infections. Altogether, our work highlights the complex dynamics of host-virus interactions during virus production with antiviral consequences for released virus particles, infected cells, and bystander cells.

Our data reveal that a major consequence of NAI is aggregation of released virions. While NA activity is thought to be essential for escape from the surface of infected cells (7, 50, 51), we find that most particles are released even when NA is inhibited by oseltamivir (Fig. 4, S5B). Instead, the dominant effect of NAI is that released particles are aggregated. Changes in the higher-order organization of particles and the contribution of NAI to these effects were profiled by combining flow virometry with digital droplet PCR to quantify average ploidy of released particle populations over short intervals. These experiments further revealed that antibodies also induce aggregation upon binding to viral antigens on infected cells. While antibodies targeting M2 aggregated free virions (Fig. S2), HA stem and NA antibodies induced aggregation of budding but not free virus particles (Fig. S1, 4). These aggregates were to varying degrees NAI-induced (Fig. 4). We also identified NAI-independent aggregation by HA stem antibodies at later timepoints, and crosslinking-dependent early size shifts occurring within 15 minutes of antibody exposure (Fig. 1). Both effects might result from antibodies directly crosslinking virions during active budding. The initial effect might wane because antigen internalization reduces surface antigen levels, making further antibody-mediated crosslinking less likely.

EM further revealed that both HA stem and NA antibodies also induce the formation of larger virions, including filaments (Fig. 5). However, the larger virions contribute marginally to the overall size shift, confirming that the antibody-induced large particles are primarily aggregates (compare Fig. 5B and Fig. S6). Notably, different antibodies caused different virion morphologies, with HA stem antibodies inducing longer, more extended filaments, while NA antibodies induced slightly elongated particles. These differences might result from variation in the cell-surface distribution of target antigen or in the extent of antibody-induced NAI. The Fab-induced delayed size shift (Fig. 2) might similarly reflect filamentation as Fab fragments cannot crosslink antigen and likely have lower levels of NAI due to their smaller size (35, 39).

In addition to altering size distributions, antibodies also reduced virus particle yields. Unlike for oseltamivir, yield reduction by HA stem and NA antibodies resulted only in part from aggregation, revealing that virus production was also attenuated (Fig. 1, 4). We propose that HA or NA depletion from the infected cell surface by antibody-induced internalization (Fig. 3) might lead to a reduced frequency of budding and consequently reduce yields. On the other hand, M2 antibodies did not have this effect despite also inducing antigen internalization (Fig. S2, 3). The extent of M2 internalization might therefore be insufficient for effective depletion, or its reduction might have little impact on budding because M2 is only a minor component of virions (11). Furthermore, while both dimerization-capable scFv constructs caused size shifts, only the scFv-tagRFP-T construct reduced yields (Fig. 2), decoupling yield and size effects. This might result from subtle differences in sfGFP and tagRFP-T dimerization affinities. At budding sites with high glycoprotein density, the resulting high local concentration of bound antibody might favor dimerization even of weakly associating variants, promoting aggregation of budding virions and enhancing size shifts. Conversely, antigen crosslinking at sites lacking active assembly might lead to their internalization instead. If these sites have lower glycoprotein density, weaker dimers might not effectively dimerize and reduce yields. The sfGFP and tagRFP-T dimerization dependence on antigen density might thus impact the relative magnitude of their yield and size effects. Notably, some monomeric fragments had the opposite effect on yields, increasing rather than decreasing it (Fig. 2). We hypothesize that these smaller molecules might act as spacers, increasing the intermolecular distance between HAs on the infected-cell surface. The larger spacing might allow fewer HAs to be incorporated per particle or reduce spontaneous HA clustering and subsequent internalization, in both cases leaving more HAs available for building additional particles.

Aggregation, filamentation, and antigen internalization may have both antiviral and proviral consequences. Virus particles formed in the presence of HA stem-binding antibody showed reduced attachment in the next round of infection (Fig. 6). This represents a novel mechanism by which these antibodies interfere with productive infection, as it occurs upon encounter of budding virus particles on the infected-cell surface rather than free, mature virus particles and without additional factors such as C1q. However, it remains unclear whether antibody-induced aggregates are always disadvantageous, or whether the benefits of larger, aggregated particles can outweigh the inhibitory effects of bound antibody. If some aggregates successfully entered cells despite attenuation, delivery of multiple genomes could influence allelic persistence or genetic exchange, accelerating viral adaptation. Filamentation may also be a proviral response to antibodies during budding, as filaments can resist antibody-mediated inhibition of cell entry (42, 43). Consistent with this, inefficient infections were associated with increased filament formation (36), suggesting that these particles might reflect a proviral adaptation under attenuation. Finally, while antigen depletion via internalization may contribute to yield reduction, it may also contribute to viral immune evasion, as depletion of surface-displayed antibody might interfere with clearance of infected cells via Fc effector functions. The balance between proviral and antiviral effects of antibody-induced changes may also be context-specific. For example, certain cell types might be more susceptible to attachment and entry of aggregates. Further study is needed to define these antibody-induced molecular events in the initial and following rounds of infection, including how they depend on antibody epitope or concentration, virus strain, and cell type.

In sum, we revealed dynamic and multifaceted effects of antibodies on the yield, physical properties, and functions of virus particles released in their presence. These insights were enabled by combining high-sensitivity flow virometry and digital droplet PCR with electron microscopy. Altogether, this study highlights the importance of investigating how immune responses impact not only infection outcomes but the properties of released virus particles in relevant biological contexts.

## Acknowledgements

The electron microscopy imaging in this paper would not have been possible without the generous support of Dr. Vinod Nair and Cindi Schwartz (Electron Microscopy Unit, Research Technologies Branch, NIAID). We would also like to thank Ruiting Xu for generating the MEDI8852 scFv-sfGFP construct. This research was supported in part by the Intramural Research Program of the National Institutes of Health (NIH). The contributions of the NIH authors are considered Works of the United States Government. The findings and conclusions presented in this paper are those of the authors and do not necessarily reflect the view of the NIH or the U.S. Department of Health and Human Services. We also acknowledge support from The G. Harold and Leila Y. Mathers Charitable Foundation (T.I.). The funders had no role in study design, data collection and analysis, decision to publish, or preparation of the manuscript.

## Author Contributions

Conceptualization: A.J.W., T.I.; Methodology: A.J.W., A.S.P., T.L.; Resources: A.J.W., E.A.P.; Investigation: A.J.W., A.S.P., T.L., E.A.P.; Formal analysis: A.J.W., E.A.P.; Validation: A.S.P., E.A.P.; Visualization: A.J.W.; Writing: A.J.W., T.I.; Review & editing: A.J.W., A.S.P., T.L., E.A.P., T.I.; Project administration and supervision: T.I.; Funding acquisition: T.I.

## Materials and Methods

### Reagents

#### Cells

Madin-Darby canine kidney (MDCK)-2,6-sialyltransferase (SIAT1) cells (Sigma-Aldrich, 05071502) were propagated in DMEM (Cytiva) supplemented with 10% FBS (Atlas Biologicals) and grown at 37°C with 5% CO2 and 100% humidity. 1 mg/mL G418 sulfate (Gold Biotechnology) was included in the growth media every other passage to select for cells retaining the plasmid for α-2,6-sialyltransferase overexpression. Expi293F cells, a gift from Theodore Pierson (NIH/VRC), were propagated in Expi293 Expression Medium (Thermo Fisher) and used for antibody expression.

#### Viruses

A/Puerto Rico/8/1934 (PR8) wild-type influenza virus was passaged in MDCK-SIAT1 cells at MOI 0.002 with 1 μg/mL TPCK-trypsin (Sigma-Aldrich) in OptiMEM (Thermo Fisher). Virus stock particle counts and size distribution were determined by flow virometry, and infectious units were determined by quantifying IAV nucleoprotein-expressing cells by flow cytometry as described (36). Work as described with IAV strain A/Puerto Rico/8/1934 was approved by the Institutional Biosafety Committee (IBC) at the Biosafety Level 2 Laboratory (BSL-2; Registration #RD-23-IV-10).

#### Antibodies

Hybridomas producing anti-IAV HA antibody H36-26 IgG and NA antibodies NA2-1C1 IgG and NA2-10E10 IgG were gifts from Jonathan Yewdell (NIH/NIAID/LVD/CBS). Hybridoma producing anti-IAV M2 monoclonal antibody 65 IgG was a gift from Xavier Saelens (VIB-UGent Center for Medical Biotechnology). The expression vectors (modified pVRC8400) for MEDI8852 IgG heavy and light chains were a gift from S. C. Harrison (Harvard Medical School). The 14C2 IgG construct was previously described (36). Antibodies were produced by transient transfection of Expi293F cells with 0.4 μg heavy chain DNA and 0.6 μg light chain DNA per 3×10^6^ cells using the ExpiFectamine 293 Transfection Kit (Gibco). Expressed antibodies were collected from the cell culture supernatant 4–7 days post transfection. IgG antibodies were purified using protein G resin (Cytiva, 17061805) and concentrated and buffer-exchanged into PBS or PBS containing 10% glycerol using a 100 kDa Amicon Ultra Centrifugal Filter (Sigma-Aldrich, UFC910008).

#### Antibody fragments

MEDI8852 Fab fragments were prepared using the Pierce Fab Preparation Kit (Thermo Fisher). To prepare the MEDI8852 F(ab’)_2_ fragments, MEDI8852 IgG was first buffer exchanged into 0.1 M sodium citrate pH 3.5 using size exclusion chromatography (Cytiva, 29148721). A 3% w/w mixture of pepsin (Sigma-Aldrich, P6887-250MG) in 0.1 M sodium citrate pH 3.5 and the prepared MEDI8852 IgG was incubated with shaking at 37°C for 2 hours. The digestion was stopped by bringing it to pH 8 with 2 M Tris base. The F(ab’)_2_ fragments were concentrated using a 50 kDa Amicon Ultra Centrifugal Filter (Sigma-Aldrich, UFC905008), and buffer exchanged into PBS using size exclusion chromatography (Cytiva, 29148721). Digestion efficiency was confirmed via SDS-PAGE. MEDI8852 scFv fragment plasmids were ordered to be synthesized and expressed by Biointron (Biointron Antibody Expression Service, CHO cells).

#### Antibody constructs

The MEDI8852 scFv-sfGFP construct (see amino acid sequence below) was designed as previously published for the 14C2 M2 antibody (52). The pVRC8400 vector backbone was digested from a pVRC8400 plasmid using NheI and NotI. The MEDI8852 variable heavy (VH) and light (VL) chain fragments were synthesized by PCR from our pVRC8400-MEDI8852-VH and -VL plasmids, respectively. The sfGFP fragment was synthesized by PCR from a plasmid expressing 14C2-scGFP.3, gifted by Nileena Velappan (Los Alamos National Laboratory). These fragments were combined into the pVRC8400 vector by Gibson assembly to create the pVRC8400-MEDI8852-scFv-sfGFP-6xHis plasmid as follows: signal peptide, MEDI8852 VL, linker, sfGFP, linker, MEDI8852 VH, 6xHis tag. The protein was produced by transfection of Expi293F cells with 0.4 μg DNA per 3×10^6^ cells using the ExpiFectamine 293 Transfection Kit (Gibco) and collected from the cell culture supernatant 7 days post transfection. The protein was purified using a HisTrap excel column (Cytiva, 17371206), concentrated using a 30 kDa Amicon Ultra Centrifugal Filter (Sigma-Aldrich, UFC903008), buffer exchanged into PBS using a 10K Slide-A-Lyzer Dialysis Cassette (Thermo Fisher, 66380), and stored in PBS containing 10% glycerol. pVRC8400-MEDI8852-scFv-eGFP-6xHis, pVRC8400-MEDI8852-scFv-tagRFP-T-6xHis, and pVRC8400-MEDI8852-scFv-tagRFP-T (E205K)-6xHis were analogously designed using published fluorophore sequences (47, 53, 54) and a different signal peptide, which is not maintained in the secreted protein (see amino acid sequences below). These constructs were ordered to be synthesized and expressed by Biointron (Biointron Antibody Expression Service, CHO cells) and were then further purified by size exclusion chromatography (Cytiva, 28990944).

**MEDI8852-scFv**-<u>sfGFP</u>:

MDAMKRGLCCVLLLCGAVFVSPSAS**DIQMTQSPSSLSASVGDRVTITCRTSQSLSSYTHWYQQKPGKAPKLLIYAASSRGSGVPSRFSGSGSGTDFTLTISSLQPEDFATYYCQQSRTFGQGTKVE**SGGSTITS<u>LFTGVVPILVELDGDVNGHKFSVRGEGEGDATNGKLTLKFICTTGKLPVPWPTLVTTLTYGVQCFSRYPDHMKRHDFFKSAMPEGYVQERTISFKDDGTYKTRAEVKFEGDTLVNRIELKGIDFKEDGNILGHKLEYNFNSHNVYITADKQKNGIKANFKIRHNVEDGSVQLADHYQQNTPIGDGPVLLPDNHYLSTQSVLSKDPNEKRDHMVLLEFVTAAGITHG</u>GGSSSGT**QVQLQQSGPGLVKPSQTLSLTCAISGDSVSSYNAVWNWIRQSPSRGLEWLGRTYYRSGWYNDYAESVKSRITINPDTSKNQFSLQLNSVTPEDTAVYYCARSGHITVFGVNVDAFDMWGQGTMVTVS**HHHHHH

**MEDI8852-scFv**-<u>eGFP</u>:

MHSSALLCCLVLLTGVRA**DIQMTQSPSSLSASVGDRVTITCRTSQSLSSYTHWYQQKPGKAPKLLIYAASSRGSGVPSRFSGSGSGTDFTLTISSLQPEDFATYYCQQSRTFGQGTKVE**SGGSTITS<u>LFTGVVPILVELDGDVNGHKFSVSGEGEGDATYGKLTLKFICTTGKLPVPWPTLVTTLTYGVQCFSRYPDHMKQHDFFKSAMPEGYVQERTIFFKDDGNYKTRAEVKFEGDTLVNRIELKGIDFKEDGNILGHKLEYNYNSHNVYIMADKQKNGIKVNFKIRHNIEDGSVQLADHYQQNTPIGDGPVLLPDNHYLSTQSALSKDPNEKRDHMVLLEFVTAAGITLG</u>GGSSSGT**QVQLQQSGPGLVKPSQTLSLTCAISGDSVSSYNAVWNWIRQSPSRGLEWLGRTYYRSGWYNDYAESVKSRITINPDTSKNQFSLQLNSVTPEDTAVYYCARSGHITVFGVNVDAFDMWGQGTMVTVS**HHHHHH

**MEDI8852-scFv**-<u>tagRFP-T</u>:

MHSSALLCCLVLLTGVRA**DIQMTQSPSSLSASVGDRVTITCRTSQSLSSYTHWYQQKPGKAPKLLIYAASSRGSGVPSRFSGSGSGTDFTLTISSLQPEDFATYYCQQSRTFGQGTKVE**SGGSTITS<u>LIKENMHMKLYMEGTVNNHHFKCTSEGEGKPYEGTQTMRIKVVEGGPLPFAFDILATSFMYGSRTFINHTQGIPDFFKQSFPEGFTWERVTTYEDGGVLTATQDTSLQDGCLIYNVKIRGVNFPSNGPVMQKKTLGWEANTEMLYPADGGLEGRTDMALKLVGGGHLICNFKTTYRSKKPAKNLKMPGVYYVDHRLERIKEADKETYVEQHEVAVARYCDLPSKLGHKLNG</u>GGSSSGT**QVQLQQSGPGLVKPSQTLSLTCAISGDSVSSYNAVWNWIRQSPSRGLEWLGRTYYRSGWYNDYAESVKSRITINPDTSKNQFSLQLNSVTPEDTAVYYCARSGHITVFGVNVDAFDMWGQGTMVTVS**HHHHHH

**MEDI8852-scFv**-<u>tagRFP-T (E205K)</u>:

MHSSALLCCLVLLTGVRA**DIQMTQSPSSLSASVGDRVTITCRTSQSLSSYTHWYQQKPGKAPKLLIYAASSRGSGVPSRFSGSGSGTDFTLTISSLQPEDFATYYCQQSRTFGQGTKVE**SGGSTITS<u>LIKENMHMKLYMEGTVNNHHFKCTSEGEGKPYEGTQTMRIKVVEGGPLPFAFDILATSFMYGSRTFINHTQGIPDFFKQSFPEGFTWERVTTYEDGGVLTATQDTSLQDGCLIYNVKIRGVNFPSNGPVMQKKTLGWEANTEMLYPADGGLEGKTDMALKLVGGGHLICNFKTTYRSKKPAKNLKMPGVYYVDHRLERIKEADKETYVEQHEVAVARYCDLPSKLGHKLNG</u>GGSSSGT**QVQLQQSGPGLVKPSQTLSLTCAISGDSVSSYNAVWNWIRQSPSRGLEWLGRTYYRSGWYNDYAESVKSRITINPDTSKNQFSLQLNSVTPEDTAVYYCARSGHITVFGVNVDAFDMWGQGTMVTVS**HHHHHH

#### Fluorescently labeled antibody

Lyophilized DyLight550 (Thermo Fisher, 62262) or AlexaFluor488 (ThermoFisher, A20000) NHS ester dye was resuspended in anhydrous dimethylsulfoxide (Sigma-Aldrich). Labeling was performed with a 100:1 molar ratio of dye to H36-26 IgG or 8G5F11 IgG (Kerafast, EB0010) in 100 mM sodium bicarbonate buffer for 1 hour at room temperature, followed by passage through a G-25 or G-50 Macro SpinColumn (Harvard Apparatus) in PBS. Labeled antibody concentrations were calculated using absorbance at 280 nm and 494 or 557 nm.

### Small-scale infections

#### General

MDCK-SIAT1 cells were grown in 24-well plates at 37°C until they formed confluent monolayers. After washing twice in HBSS (Thermo Fisher), 35 μL PR8 virus diluted in OptiMEM (Thermo Fisher) was added at MOI 0.6 and incubated at room temperature for 1 hour with frequent shaking. After attachment, cells were washed twice with HBSS (Thermo Fisher) and 300 μL OptiMEM (Thermo Fisher) was added. Trypsin was omitted to inhibit spread. Infections were incubated at 34°C with 5% CO2 and 100% humidity. Samples were stored in PCR tubes (Thermo Fisher, 14-230-210) at −80°C until analysis.

#### Infections for transient antibody treatment

At 21 h.p.i., cells were washed twice with HBSS (Thermo Fisher), and media was replaced with 300 μL OptiMEM (Thermo Fisher) containing the indicated concentrations of antibody. 20-μL supernatant samples were collected at 15, 30, 60, 90, 120 minutes post-treatment and analyzed by flow virometry.

#### Infections with bacterial neuraminidase pretreatment

At 20 h.p.i., bacterial neuraminidase (bNA; neuraminidase from *Vibrio cholerae*, Sigma-Aldrich, N7885-2UN) was added to the media at a final dilution of 1:1000 and incubated for 2 hours at 34°C. The media was then replaced with OptiMEM (Thermo Fisher) containing bNA as in the pretreatment and 1000 nM oseltamivir (LGC Standards, TRC-O700980-25MG), 2000 nM MEDI8852 IgG, or 500 nM NA2-1C1 IgG. Supernatants were harvested at 24 h.p.i. and analyzed by flow virometry and digital droplet PCR. Mock controls without bNA pretreatment were included.

#### Infections for immunofluorescence

At 21 h.p.i., cells were washed twice with HBSS (Thermo Fisher) and media was replaced with 300 μL OptiMEM (Thermo Fisher) containing 500 nM MEDI8852 IgG, NA2-1C1 IgG, or NA2-10E10 IgG. At 120 minutes post-treatment, cells were washed thrice with PBS, fixed with 4% PFA (Fisher, AAJ61899AP) in PBS for 30 minutes, and washed thrice with PBS again. Cells were analyzed by immunofluorescence.

#### Infections for binding assay

At 20 h.p.i., cells were washed twice with HBSS (Thermo Fisher) and media was replaced with 300 μL OptiMEM (Thermo Fisher) with or without 2000 nM MEDI8852 IgG. The supernatant was collected at 24 h.p.i. and used as the input virus in the binding assay.

### Infections for electron microscopy

MDCK-SIAT1 cells were grown in 175 cm^2^ flasks at 37°C until they formed confluent monolayers. After washing twice with HBSS (Thermo Fisher), 3 mL PR8 virus diluted in OptiMEM (Thermo Fisher) was added at MOI 0.6 and incubated at room temperature for 1 hour with frequent shaking. After attachment, cells were washed twice with HBSS (Thermo Fisher) and 20 mL OptiMEM (Thermo Fisher) was added. Trypsin was omitted to inhibit spread. After 4 hours at 34°C, the supernatant was removed and replaced with fresh media alone, media containing 500 nM MEDI8852 IgG, or media containing 500 nM NA2-1C1 IgG. After another 20 hours of incubation, the supernatant was collected and analyzed by flow virometry. For electron microscopy, 1 flask was used for each mock condition, 3 flasks were pooled for the MEDI8852 IgG condition, and 2 flasks were pooled for the NA2-1C1 IgG condition. The supernatant was clarified by centrifugation at 1000 xg for 10 min, treated with a 1:5000 dilution of bNA (Sigma-Aldrich, N7885-2UN), incubated at 34°C for 1 hour, and returned to ice. The sample was then adjusted to 30 mL with PBS as necessary, layered onto 2.5 mL 60% sucrose and 5 mL 20% sucrose, and spun at 100000 xg for 1 hour at 4°C. The band at the sucrose layer interface was collected. The band from the MEDI8852 IgG condition was fixed by adding 37% formaldehyde to a final concentration of 4%, incubating for 15 minutes at RT, and diluting to 14 mL in PBS. 14 mL of the fixed virus from the MEDI8852 IgG condition or the unfixed bands from the mock and NA2-1C1 IgG conditions was layered onto 800 μL 60% sucrose and 2 mL 20% sucrose and spun at 100000 xg for 1 hour at 4°C. The 60% sucrose layer and virus interface were collected and diluted 1:1 in PBS. The sample diluted to 2.2 mL with PBS, layered onto 60 μL 60% sucrose, and spun at 100000 xg for 90 minutes at 4°C. The purified, concentrated, bNA-treated virus was analyzed by electron microscopy.

### Flow virometry

#### General

DyLight550-labeled H36-26 IgG stock solution (see *Fluorescently labeled antibody* above) was diluted to 50 nM in 0.2% BSA and HNE20 (20 mM HEPES NaOH pH 7.4, 150 mM NaCl and 0.2 mM EDTA). Infected-cell supernatants were left undiluted or diluted up to 1:10 in HNE20 and combined 1:1 with antibody dilution in BSA. Binding reactions were incubated at room temperature for 1 hour, then diluted 1:250 in HNE20. Flow virometry was performed using the CytoFLEX S platform (Beckman Coulter). Laser powers were 70 mW for violet and 50 mW for yellow. Gain values were set to 250 for VSSC and 1,000 for RFP. Samples were triggered on violet side-scatter area (3,500 a.u. threshold) and acquired for 600 seconds or until acquiring 25,000 events in the virus particle gate (Fig. 1A). Analysis was performed in Cytexpert 2.5.

#### Aggregation testing

Supernatants collected from PR8 infections immediately prior to treatment at 21 hours post-infection (see *Transient antibody treatment infections* above) were diluted to 7e5 particles/µL or 4e6 particles/µL (resembling the virus concentrations at 15- and 120- minutes of mock treatment, respectively) in the highest concentration of each antibody treatment (2000 nM MEDI8852 IgG and F(ab’)_2_, 4000 nM MEDI8852 Fab and scFv, 8000 nM MEDI8852 scFv constructs, and 500 nM NA and M2 antibodies) in OptiMEM (Thermo Fisher). Samples were incubated at 34°C for 15 or 120 minutes, respectively, and analyzed by flow virometry.

#### Antibody fragment binding testing

Samples containing 1e6 virus particles/µL from mock-treated PR8 infections (see **Small-scale infections** above), 25 nM DyLight550-labeled H36-26 IgG, and 100 nM MEDI8852 scFv-eGFP were prepared in 0.2% BSA and HNE20, with or without 500 nM MEDI8852 Fab fragment or 500 nM MEDI8852 scFv fragment. Samples were incubated at room temperature for 1 hour, diluted 1:250 in HNE20, and analyzed by flow virometry.

#### Deaggregation testing

Supernatants from oseltamivir- and antibody-treated PR8 infections with mock pretreatment of cells (see *Bacterial neuraminidase infections* above) were analyzed by flow virometry immediately after collection and after incubating at 37°C for 1 hour in bNA (Sigma-Aldrich, N7885-2UN). bNA was diluted in OptiMEM (Thermo Fisher) and used at a final dilution of 1:1000.

#### Binding assay

35 µL supernatant from mock- or MEDI8852-treated PR8 infections (see *Infections for binding assay* above) was added to MDCK cells and incubated at room temperature. Unbound virus was collected after 1 hour. The input virus and unbound virus were analyzed by flow virometry.

#### Titering VSV stock

AlexaFluor488-labeled 8G5F11 IgG stock solution (see *Fluorescently labeled antibody* above) was diluted 1:200 in PBS. VSV stock of the Indianna serotype, a gift from Jonathan Yewdell and Ivan Kosik (NIH/NIAID/LVD/CBS), was diluted 1:10 in PBS. The antibody and virus dilutions were combined 1:1 and binding reactions were incubated at room temperature for 1 hour, then diluted 1:250 in PBS. The 1:400 total dilution of antibody produced the best separation between noise and VSV signal, as identified by titration.

### Confocal immunofluorescence microscopy

Coverglasses were sterilized with 70% ethanol, treated with 0.01% poly-L lysine for 30 minutes at room temperature, washed twice with water, and washed once with HBSS. Cell seeding, infection, treatment, and fixing were performed as described (see *Immunofluorescence infections* above). Cells were then permeabilized and stained with 300 nM DAPI and 1 µg/mL AF488-goat-anti-mouse secondary antibody (Invitrogen, A11001) in PBS containing 3% BSA and 0.25% TritonX-100 at room temperature for 1 hour in the dark. Cells were washed thrice with PBS and stained with 2 µg/mL CellMask Blue (Thermo Fisher, H32720) in PBS at room temperature for 1 hour. Cells were washed thrice with PBS, mounted on slides with Prolong anti-fade (Invitrogen, P36961), and cured for 24 hours at room temperature in the dark. Samples were imaged at the Biological Imaging Section (NIH/NIAID/RTB) on a Leica SP8 (690) DMI6000 confocal microscope (Leica Microsystems) using an HC PL APO CS2×40/1.30 oil objective. Imaging and image processing were performed in LAS X and Fiji, respectively.

### Digital droplet PCR

130 µL virus supernatant from PR8 infections (see *Bacterial neuraminidase infections* above) was mixed with 10 µL VSV stock of known concentration (see *Titering VSV stock* above). To remove extracellular viral RNA not enclosed within virions, RNase A was added to a final concentration of 50 ng/mL and the mixture was incubated at 37 °C for 30 minutes. Viral RNA was extracted using the QIAamp Viral RNA Mini Kit (Qiagen) and used as the template for reverse transcription with M-MuLV Reverse Transcriptase (New England Biolabs) using primers specific for influenza A virus and VSV (Uni-12, 5′-ACGCGTGATCAGCAAAAGCAGG-3′; VSV-Tr-F, 5′-AAATCATGAGGAGACTCCAAAC-3′). Genome copies of VSV and IAV were quantified using the QX200 Droplet Digital PCR System (Bio-Rad) with the following primer sets: VSV-Tr-F, 5′-AAATCATGAGGAGACTCCAAAC-3′; VSV-Tr-R, 5′-ACGAAGACCACAAAACCAG-3′; PB2-all-F, 5′-AATCTAATGTCGCAGTCTCG-3′; and PB2-all-R, 5′-TCATCCTAAGTGCTGGGTTC-3′. The raw genome-to-particle ratio was calculated from the genome copies obtained by ddPCR and the particle count obtained by flow virometry. To adjust for gains or losses during processing, the raw genome-to-particle ratio for IAV was divided by the genome-to-particle ratio of VSV used as an internal control, yielding an adjusted genome-to-particle ratio. The values were further normalized to those of mock-pretreated, mock-treated samples to facilitate comparison between experiments.

### Electron microscopy

40 µL virus (see **Large-scale infections** above) was treated with 1 µL rimantadine, diluted to 1 mM in PBS (Sigma-Aldrich, PHR3267-500MG), to a final rimantidine concentration of 25 µM. Rimantadine inhibits the M2 proton channel on virions and might reduce fracturing/disruption of virions during EM preparation (45). If samples were diluted before making grids, they were diluted in PBS with 25 µM rimantadine. To prepare the grids, Carbon Film, 400 Mesh Copper grids (Electron Microscopy Sciences, CF400-CU-50) were glow discharged for 30 seconds at 1.5 mA in 39 mBar air using a PELCO easiGlow Glow Discharge Cleaning System (Ted Pella, 91000). 5 µL sample was applied to the grid and incubated for 120 seconds. The sample was wicked off, rinsed briefly in PTA (Ted Pella, 19402) dissolved to 2% in PBS and brought to pH 7.5 with NaOH, and stained in 2% PTA pH 7.5 for 30 seconds. The PTA was then wicked off, and the grid was dried with gentle vacuum. Grids were imaged in a Talos (ThermoFisher Scientific) transmission electron microscope operating at 120 kV with a Ceta (ThermoFisher Scientific) camera. Automated data collection was performed using SerialEM (University of Colorado, Version 4.2.12). A grid square was chosen manually at low magnification and an array of points was added with a spacing of 1.5 µm. Motion corrected images were then automatically acquired at each point with a defocus set to -1 µm at a nominal magnification of 28000x (0.51 nm pixel).

### Statistics and reproducibility

Flow virometry experiments were not randomized and the investigators were not blinded to allocation during experiments and outcome assessment. Analysis was automated and independent of human interpretation. For example, gates in flow virometry were set using a control and applied to all samples. Yield and size measurements <1,000 particles/μL, where noise measured in negative samples reached ∼10% of the virion count, were excluded from analysis. Additionally, one data point in Fig. S2B (14C2) was identified as an outlier and excluded based on Grubb’s test with a significance level of α=0.5. Electron microscopy was performed by a biologist in the Rocky Mountain Laboratories Research Technologies Section. Unaware of the sample identities, she identified a grid square with good stain, verified the quality of the sample/stain, defined a large imaging area spanning the grid square, then started an automated acquisition. Images were scrambled and given non-identifying names before particle length measurements, which were performed blind to the treatment condition. The number of particles imaged was limited by the lowest-concentration samples. One biological replicate was imaged after independent confirmation that it exhibited the expected treatment response by flow virometry (see Fig. S6). For all experiments, data distribution was assumed to be normal, but this was not formally tested.

## References

1. Choppin PW, Murphy JS, Tamm I. 1960. Studies of two kinds of virus particles which comprise influenza A2 virus strains. III. Morphological characteristics: independence to morphological and functional traits. J Exp Med 112:945–952.

2. Noda T. 2012. Native morphology of influenza virions. Front Microbiol 2:269.

3. Leser GP, Lamb RA. 2005. Influenza virus assembly and budding in raft-derived microdomains: A quantitative analysis of the surface distribution of HA, NA and M2 proteins. Virology 342:215–227.

4. Dahmani I, Ludwig K, Chiantia S. 2019. Influenza A matrix protein M1 induces lipid membrane deformation via protein multimerization. Biosci Rep 39: BSR20191024.

5. Zhang J, Lamb RA. 1996. Characterization of the membrane association of the influenza virus matrix protein in living cells. Virology 225:255–266.

6. Rossman JS, Jing X, Leser GP, Lamb RA. 2010. Influenza virus M2 protein mediates ESCRT-independent membrane scission. Cell 142:902–913.

7. Palese P, Tobita K, Ueda M, Compans RW. 1974. Characterization of temperature-sensitive influenza virus mutants defective in neuraminidase. Virology 61:397–410.

8. Lamb RA, Zebedee SL, Richardson CD. 1985. Influenza virus M2 protein is an integral membrane protein expressed on the infected-cell surface. Cell 40:627–633.

9. Mora R, Rodriguez-Boulan E, Palese P, García-Sastre A. 2002. Apical budding of a recombinant influenza A virus expressing a hemagglutinin protein with a basolateral localization signal. J Virol 76:3544–3553.

10. Harris A, Cardone G, Winkler DC, Heymann JB, Brecher M, White JM, Steven AC. 2006. Influenza virus pleiomorphy characterized by cryoelectron tomography. Proc Natl Acad Sci U S A 103:19123–27.

11. Zebedee SL, Lamb RA. 1988. Influenza A virus M2 protein: monoclonal antibody restriction of virus growth and detection of M2 in virions. J Virol 62:2762–2772.

12. Brandenburg B, Koudstaal W, Goudsmit J, Klaren V, Tang C, Bujny MV, Korse HJWM, Kwaks T, Otterstrom JJ, Juraszek J, van Oijen AM, Vogels R, Friesen RHE. 2013. Mechanisms of hemagglutinin targeted influenza virus neutralization. PLoS One 8:e80034.

13. Bizebard T, Gigant B, Rigolet P, Rasmussen B, Diat O, Bösecke P, Wharton SA, Skehel JJ, Knossow M. 1995. Structure of influenza virus haemagglutinin complexed with a neutralizing antibody. Nature 376:92–94.

14. Knossow M, Gaudier M, Douglas A, Barrère B, Bizebard T, Barbey C, Gigant B, Skehel JJ. 2002. Mechanism of neutralization of influenza virus infectivity by antibodies. Virology 302:294–298.

15. Corti D, Voss J, Gamblin SJ, Codoni G, Macagno A, Jarrossay D, Vachieri SG, Pinna D, Minola A, Vanzetta F, Silacci C, Fernandez-Rodriguez BM, Agatic G, Bianchi S, Giacchetto-Sasselli I, Calder L, Sallusto F, Collins P, Haire LF, Temperton N, Langedijk JPM, Skehel JJ, Lanzavecchia A. 2011. A neutralizing antibody selected from plasma cells that binds to group 1 and group 2 influenza A hemagglutinins. Science 333:850–856.

16. Ekiert DC, Bhabha G, Elsliger MA, Friesen RHE, Jongeneelen M, Throsby M, Goudsmit J, Wilson IA. 2009. Antibody recognition of a highly conserved influenza virus epitope. Science 324:246–251.

17. Ekiert DC, Friesen RHE, Bhabba G, Kwaks T, Jongeneelen M, Yu W, Ophorst C, Cox F, Korse HJWM, Brandenburg B, Vogels R, Brakenhoff JPJ, Kompier R, Koldijk MH, Cornelissen LAHM, Poon LLM, Peiris M, Koudstaal W, Wilson IA, Goudsmit J. 2011. A highly conserved neutralizing epitope on group 2 influenza A viruses. Science 333:843–850.

18. Okuno Y, Isegawa Y, Sasao F, Ueda S. 1993. A common neutralizing epitope conserved between the hemagglutinins of influenza A virus H1 and H2 strains. J Virol 67:2552–2558.

19. Sui J, Hwang WC, Perez S, Wei G, Aird D, Chen L, Santelli E, Stec B, Cadwell G, Ali M, Wan H, Murakami A, Yammanuru A, Han T, Cox NJ, Bankston LA, Donis RO, Liddington RC, Marasco WA. 2009. Structural and functional bases for broad-spectrum neutralization of avian and human influenza A viruses. Nat Struct Mol Biol 16:265–273.

20. Throsby M, van den Brink E, Jongeneelen M, Poon LLM, Alard P, Cornelissen L, Bakker A, Cox F, van Deventer E, Guan Y, Cinatl J, ter Meulen J, Lasters I, Carsetti R, Peiris M, de Kruif J, Goudsmit J. 2008. Heterosubtypic neutralizing monoclonal antibodies cross-protective against H5N1 and H1N1 recovered from human IgM+ memory B cells. PLoS One 3:e3942.

21. Williams JA, Gui L, Hom N, Mileant A, Lee KK. 2018. Dissection of epitope-specific mechanisms of neutralization of influenza virus by intact IgG and Fab fragments. J Virol 92:e02006–17.

22. Yamauchi K, Muramoto Y, Fujita-Fujiharu Y, Hirabayashi A, Nogami C, Ishida T, Sugita Y, Saito S, Sano K, Suzuki T, Nakano M, Noda T. 2026. Multimerization of secretory IgA enhances antiviral activity by aggregating influenza A virus particles. Commun Biol 10.1038/s42003-026-09783-9

23. Kosik I, Da Silva Santos J, Angel M, Hu Z, Holly J, Gibbs JS, Gill T, Kosikova M, Li T, Bakhache W, Dolan PT, Xie H, Andrews SF, Gillespie RA, Kanekiyo M, McDermott AB, Pierson TC, Yewdell JW. 2024. C1q enables influenza hemagglutinin stem binding antibodies to block viral attachment and broadens the antibody escape repertoire. Sci Immunol 9:eadj9534.

24. DiLillo DJ, Tan GS, Palese P, Ravetech JV. 2014. Broadly neutralizing hemagglutinin stalk–specific antibodies require FcγR interactions for protection against influenza virus *in vivo*. Nat Med 20:143–151.

25. El Bakkouri K, Descamps F, De Filette M, Smet A, Festjens E, Birkett A, Van Rooijen N, Verbeek S, Fiers W, Saelens X. 2011. Universal vaccine based on ectodomain of matrix protein 2 of influenza A: Fc receptors and alveolar macrophages mediate protection. J Immunol 186:1022–1031.

26. Huber VC, Lynch JM, Bucher DJ, Le J, Metzger DW. 2001. Fc receptor-mediated phagocytosis makes a significant contribution to clearance of influenza virus infections. J Immunol 166:7381–7388.

27. Kopf M, Abel B, Gallimore A, Carroll M, Bachmann MF. 2002. Complement component C3 promotes T-cell priming and lung migration to control acute influenza virus infection. Nat Med 8:373–378.

28. Lei R, Kim W, Lv H, Mou Z, Scherm MJ, Schmitz AJ, Turner JS, Tan TJC, Wang Y, Ouyang WO, Liang W, Rivera-Cardona J, Teo C, Graham CS, Brooke CB, Presti RM, Mok CKP, Krammer F, Dai X, Ellebedy AH, Wu NC. 2023. Leveraging vaccination-induced protective antibodies to define conserved epitopes on influenza N2 neuraminidase. Immunity 56:2621–2634.

29. Mullarkey CE, Bailey MJ, Golubeva DA, Tan GS, Nachbagauer R, He W, Novakowski KE, Bowdish DM, Miller MS, Palese P. 2016. Broadly neutralizing hemagglutinin stalk-specific antibodies induce potent phagocytosis of immune complexes by neutrophils in an Fc-dependent manner. mBio 7:e01624–16.

30. O’Brien KB, Morrison TE, Dundore DY, Heise MT, Schultz-Cherry S. 2011. A protective role for complement C3 protein during pandemic 2009 H1N1 and H5N1 influenza A virus infection. PLoS One 6:e17377.

31. Van den Hoecke S, Ehrhardt K, Kolpe A, El Bakkouri K, Deng L, Grootaert H, Schoonooghe S, Smet A, Bentahir M, Roose K, Schotsaert M, Schepens B, Callewaert N, Nimmerjahn F, Staeheli P, Hengel H, Saelens X. 2017. Hierarchical and redundant roles of activating FcγRs in protection against influenza disease by M2e-specific IgG1 and IgG2a antibodies. J Virol 91:e02500–16.

32. He Y, Guo Z, Subiaur S, Benegal A, Vahey MD. 2024. Antibody inhibition of influenza A virus assembly and release. J Virol 98:e01398–23.

33. Hughey PG, Roberts PC, Holsinger LJ, Zebedee SL, Lamb RA, Compans RW. 1995. Effects of antibody to the influenza A virus M2 protein on M2 surface expression and virus assembly. Virology 212:411–421.

34. Kolpe A, Arista-Romero M, Schepens B, Pujals S, Saelens X, Albertazzi L. 2019. Super-resolution microscopy reveals significant impact of M2especific monoclonal antibodies on influenza A virus filament formation at the host cell surface. Sci Rep 9:4450.

35. Kosik I, Angeletti D, Gibbs JS, Angel M, Takeda K, Kosikova M, Nair V, Hickman HD, Xie H, Brooke CB, Yewdell JW. 2019. Neuraminidase inhibition contributes to influenza A virus neutralization by anti-hemagglutinin stem antibodies. J Exp Med 216:304–316.

36. Partlow EA, Jaeggi-Wong A, Planitzer SD, Berg N, Li Z, Ivanovic T. 2025. Influenza A virus rapidly adapts particle shape to environmental pressures. Nat Microbiol 10:784–794.

37. Yamayoshi S, Uraki R, Ito M, Kiso M, Nakatsu S, Yasuhara A, Oishi K, Sasaki T, Ikuta K, Kawaoka Y. 2017. A broadly reactive human anti-hemagglutinin stem monoclonal antibody that inhibits influenza A virus particle release. EBioMedicine 17:182–191.

38. Stadlbauer D, Zhu X, McMahon M, Turner JS, Wohlbold TJ, Schmitz AJ, Strohmeier S, Yu W, Nachbagauer R, Mudd PA, Wilson IA, Ellebedy AH, Krammer F. 2019. Broadly protective human antibodies that target the active site of influenza virus neuraminidase. Science 366:499–504.

39. Chen YQ, Lan LY-L, Huang M, Henry C, Wilson PC. 2019. Hemagglutinin stalkreactive antibodies interfere with influenza virus neuraminidase activity by steric hindrance. J Virol 93:e01526–18.

40. Kosik I, Yewdell JW. 2017. Influenza A virus hemagglutinin specific antibodies interfere with virion neuraminidase activity via two distinct mechanisms. Virology 500:178–183.

41. Vahey MD, Fletcher DA. 2019. Low-fidelity assembly of influenza A virus promotes escape from host cells. Cell 176:281–294.

42. Li T, Li Z, Deans EE, Mittler E, Liu M, Chandran K, Ivanovic T. 2021. The shape of pleomorphic virions determines resistance to cell-entry pressure. Nat Microbiol 6:617–629.

43. Peterl S, Lahr CM, Schneider CN, Meyer J, Podlipensky X, Lechner V, Villiou M, Eis L, Klein S, Funaya C, Cavalcanti-Adam EA, Graw F, Selhuber-Unkel C, Rohr K, Chlanda P. 2025. Morphology-dependent entry kinetics and spread of influenza A virus. EMBO J 44:3959–3982.

44. Noda T, Sagara H, Yen A, Takada A, Kida H, Cheng RH, Kawaoka Y. 2006. Architecture of ribonucleoprotein complexes in influenza A virus particles. Nature 439:490–492.

45. Rossman JS, Leser GP, Lamb RA. 2012. Filamentous influenza virus enters cells via macropinocytosis. J Virol 86:10950–10960.

46. Scott DJ, Gunn NJ, Yong KJ, Wimmer VC, Veldhuis NA, Challis LM, Haidar M, Petrou S, Bathgate RAD, Griffin MDW. 2018. A Novel Ultra-Stable, Monomeric Green Fluorescent Protein For Direct Volumetric Imaging of Whole Organs Using CLARITY. Sci Rep 8:667.

47. Beyrent E, Wei DT, Beacham, GM, Park S, Zheng J, Paszek MJ, Hollopeter G. 2024. Dimerization activates the Inversin complex in *C. elegans*. Mol Biol Cell 35:ar127,1–12.

48. Pahara J, Shi H, Chen X, Wang Z. 2010. Dimerization drives PDGF receptor endocytosis through a C-terminal hydrophobic motif shared by EGF receptor. Exp Cell Res 316:2237–2250.

49. Wang Q, Villeneuve G, Wang Z. 2005. Control of epidermal growth factor receptor endocytosis by receptor dimerization, rather than receptor kinase activation. EMBO Rep 6:942–948.

50. Mori K, Murano K, Ohniwa RL, Kawaguchi A, Nagata K. 2015. Oseltamivir expands quasispecies of influenza virus through cell-to-cell transmission. Sci Rep 5:9163.

51. Palese P, Compans RW. 1976. Inhibition of influenza virus replication in tissue culture by 2-deoxy-2,3-dehydro-N-trifluoroacetylneuraminic acid (FANA): Mechanism of action. J Gen Vir 33:159–163.

52. Velappan N, Micheva-Viteva S, Adikari SH, Waldo GS, Lillo AM, Bradbury ARM. 2020. Selection and verification of antibodies against the cytoplasmic domain of M2 of influenza, a transmembrane protein. mAbs 12:e1843754.

53. Cormack BP, Valdivia RH, Falkow S. 1996. FACS-optimized mutants of the green fluorescent protein (GFP). Gene 173:33–38.

54. Shaner NC, Lin MZ, McKeown MR, Steinbach PA, Hazelwood KL, Davidson MW, Tsien RY. 2008. Improving the photostability of bright monomeric orange and red fluorescent proteins. Nat Methods 5:545–551.

