## Supplementary figures and images for "Antibodies to influenza A virus hemagglutinin and neuraminidase limit egress and alter the physical properties of released virus particles"

### Figure S1

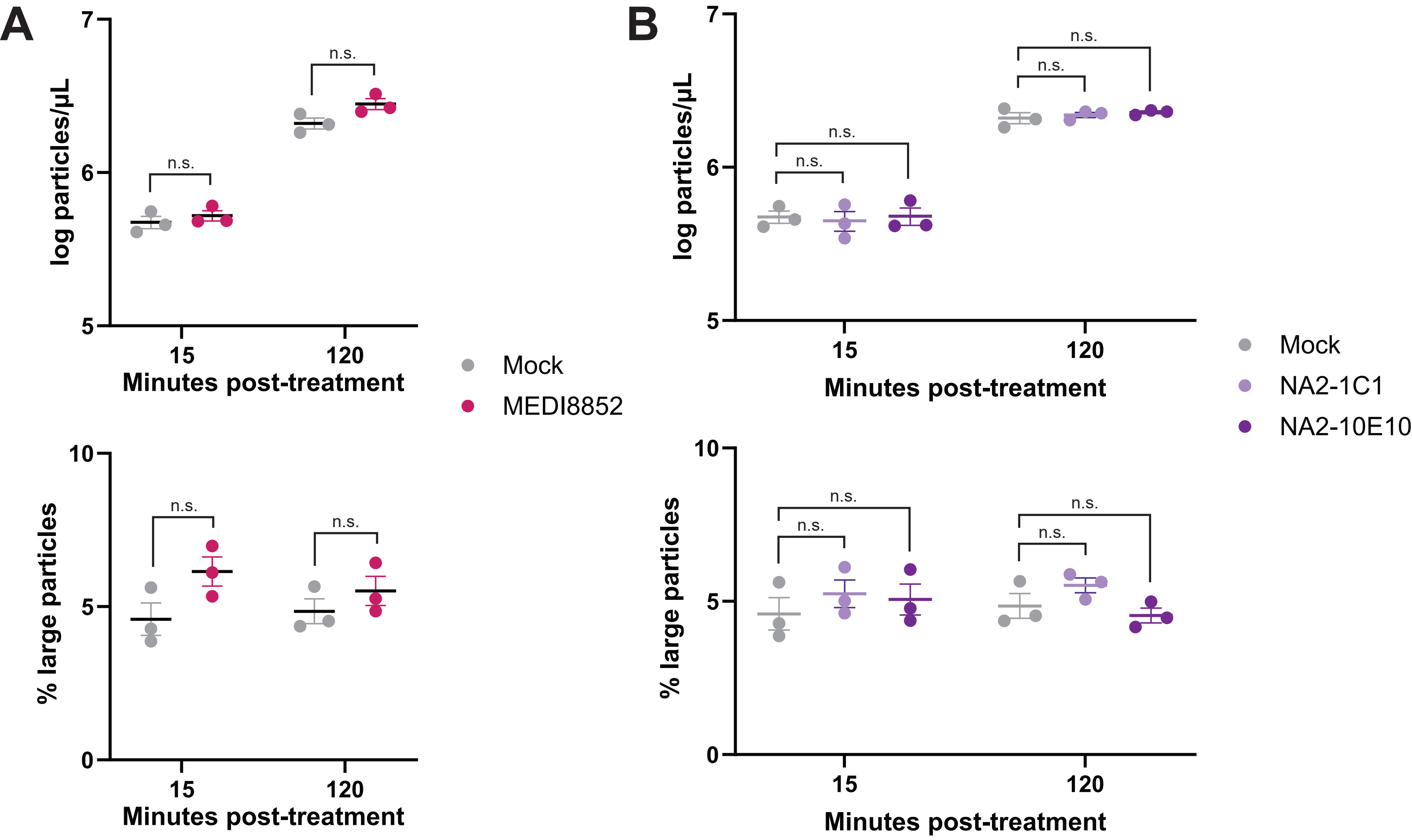

### Figure S2

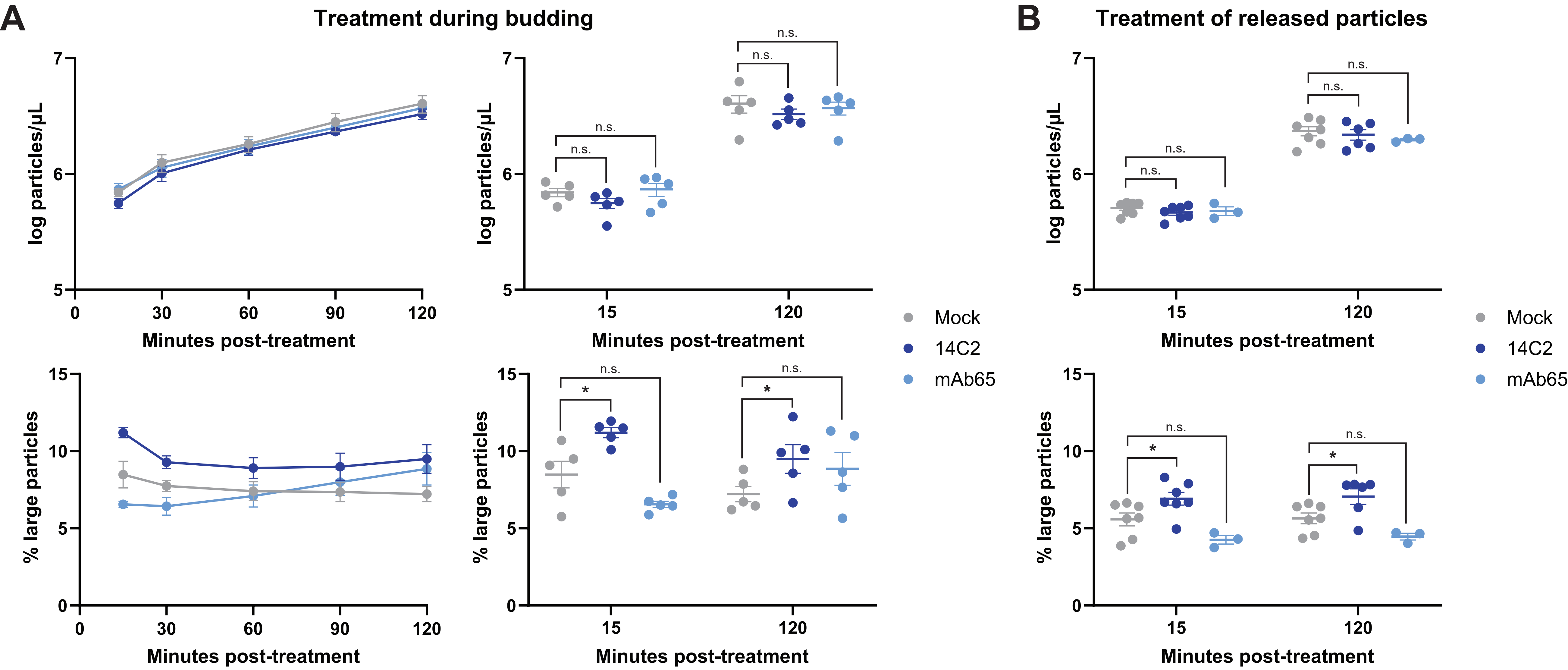

### Figure S3

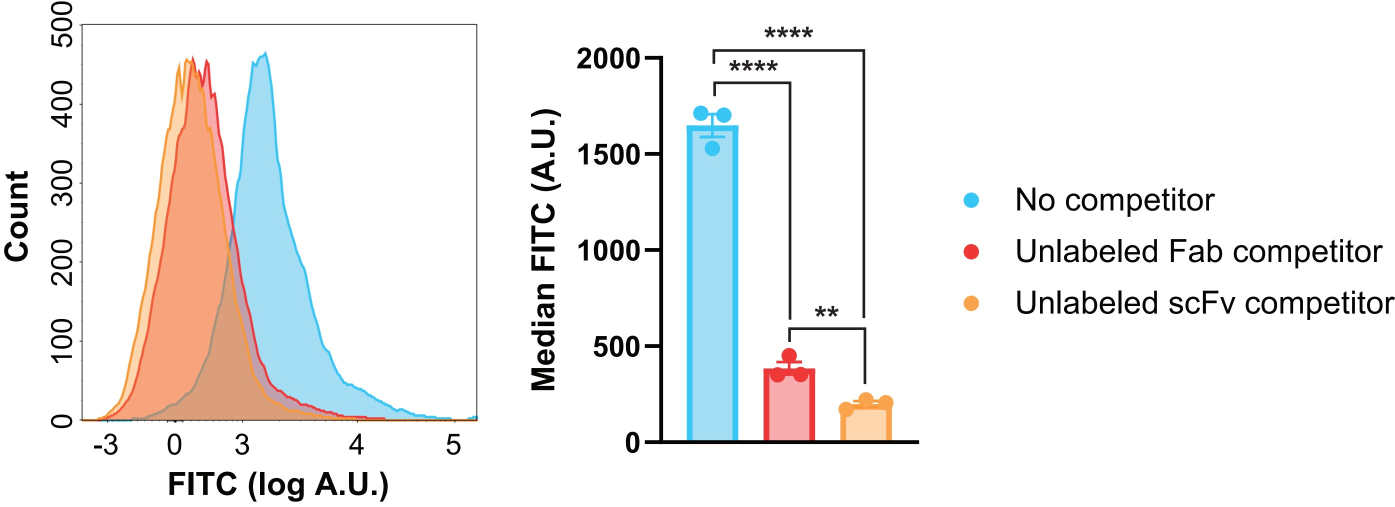

### Figure S4

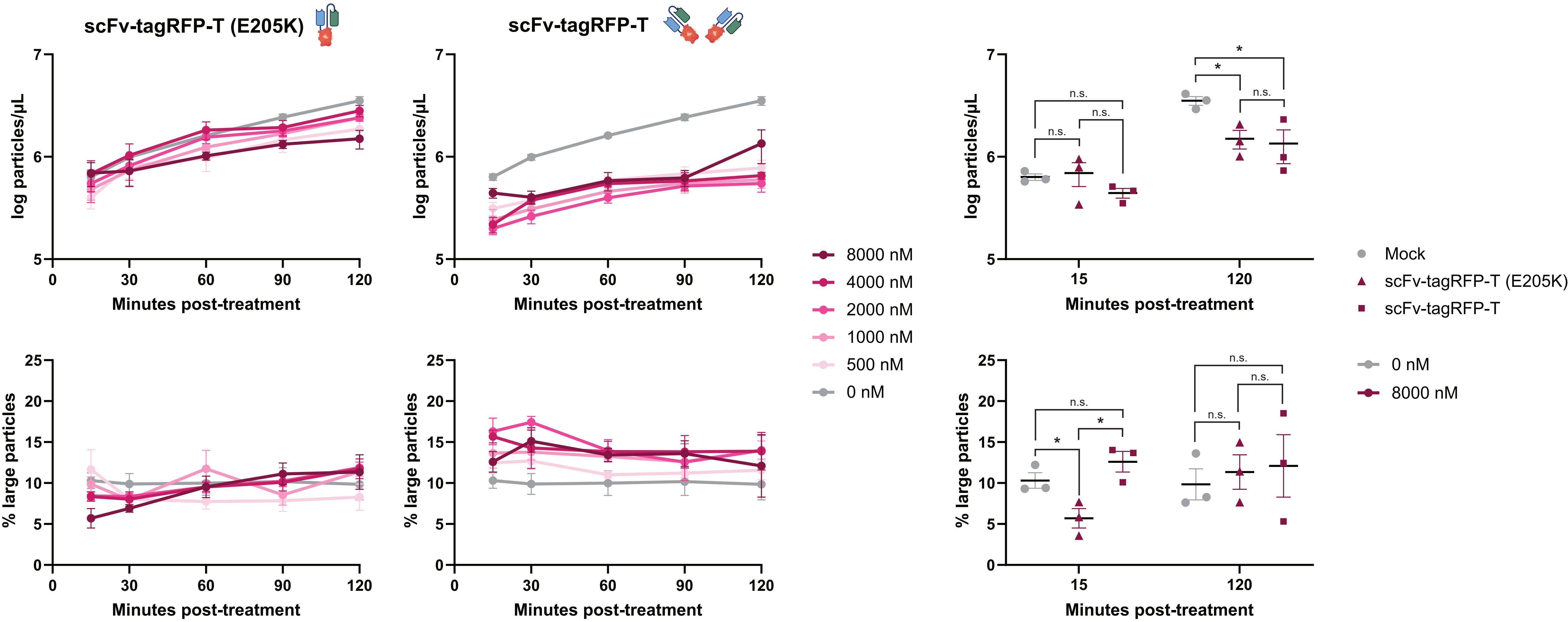

### Figure S5

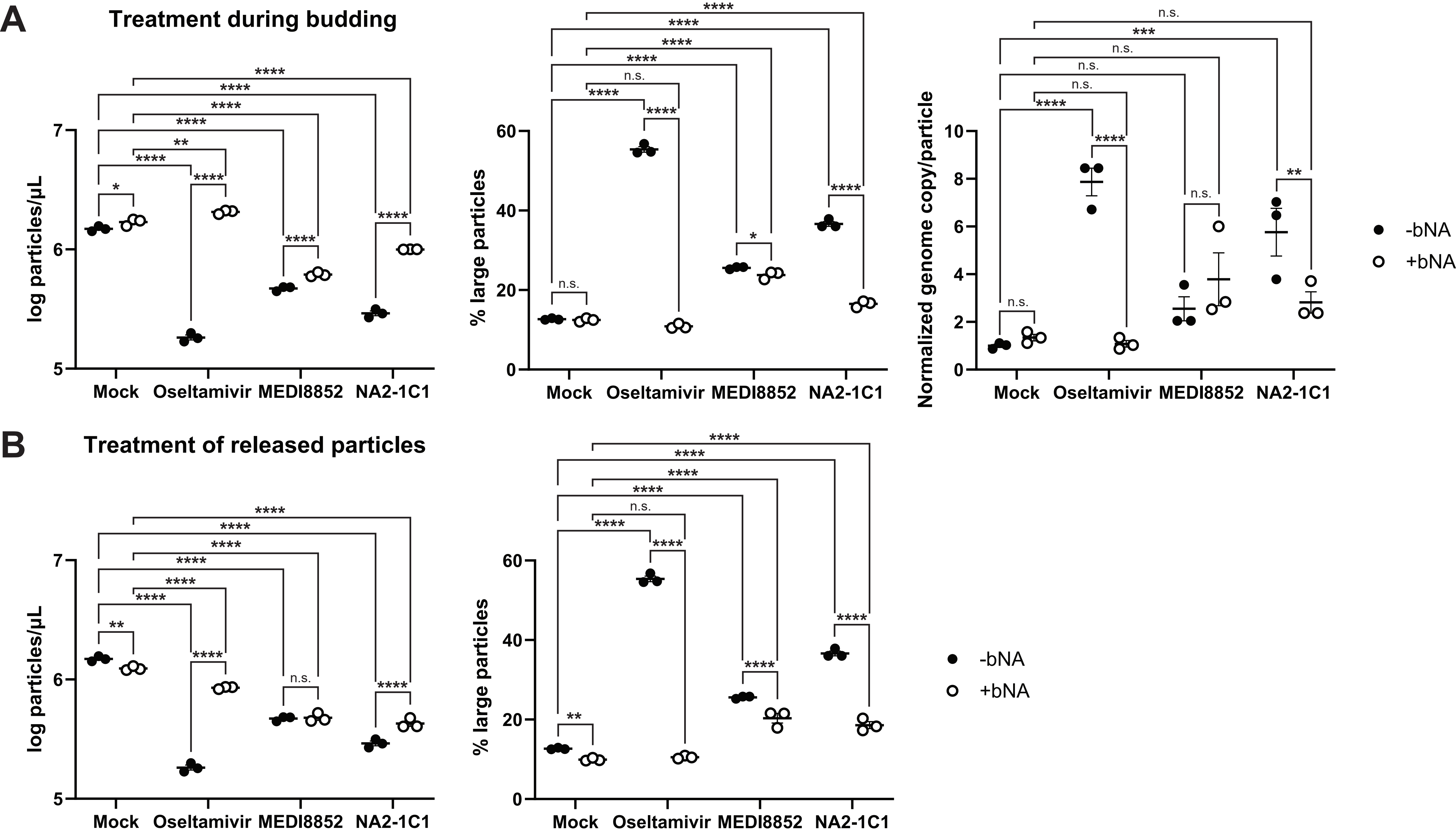

### Figure S6

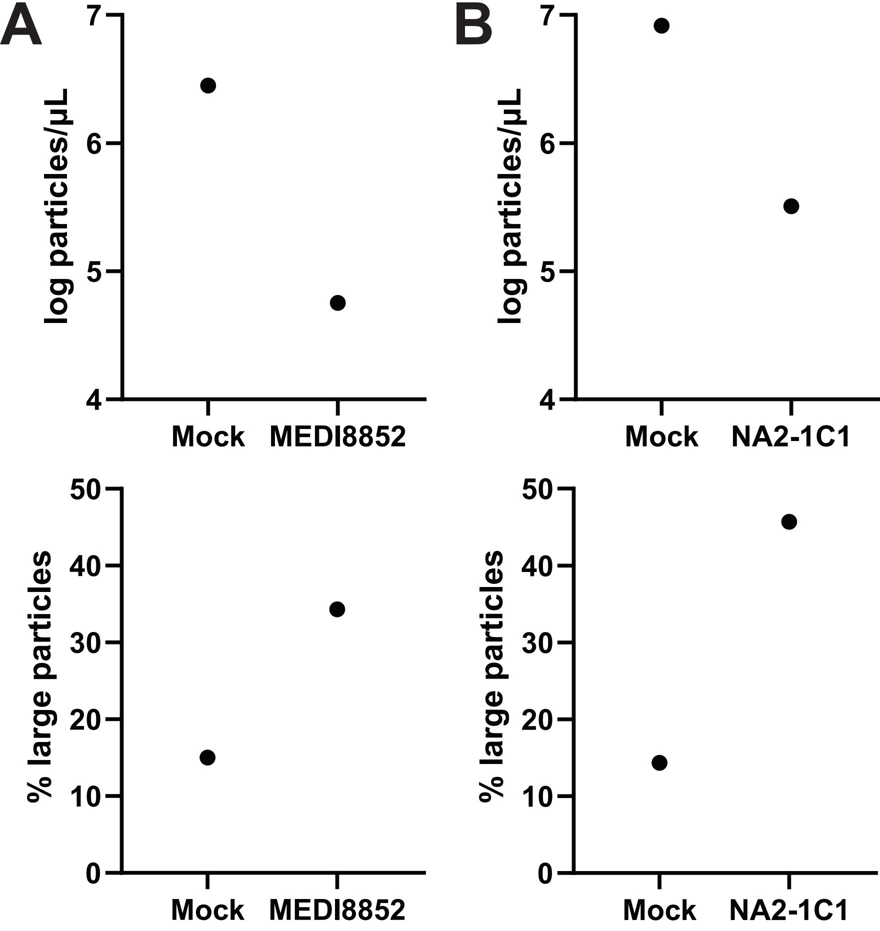
